# Proteomics-based characterization of ribosome heterogeneity in adult mouse organs

**DOI:** 10.1101/2024.02.23.581691

**Authors:** Marie R Brunchault, Anne-Marie Hesse, Julia Schaeffer, Charlotte Decourt, Florence Combes, Homaira Nawabi, Yohann Couté, Stephane Belin

**Author notes:** These authors share co-first authorship and are listed by alphabetical order (Marie R Brunchault, Anne-Marie Hesse, Julia Schaeffer).

## Abstract

While long thought to be invariable in all cellular organisms, evidence has emerged that the translation process, *i.e.* protein assembly from mRNA sequence decoding, is regulated by variable features of the translation machinery. Notably, ribosomes, the functional units of protein synthesis, display variations in their composition, depending on the developmental stage, cell type or physiopathological context, thus providing a new level of actionable regulation of gene expression. Yet, a comprehensive map of the heterogeneity of ribosome composition in ribosomal proteins (RPs) in different organs and tissues is not available. In this work, we explored tissue-specific ribosome heterogeneity using mass spectrometry-based quantitative proteomic characterization of ribosomal fractions purified from 14 adult mouse organs and tissues. We performed crossed clustering and statistical analyses of RP composition to highlight stable, variable and tissue-specific RPs across organs and tissues. Focusing on specific RPs, we validated their relative abundance with a targeted proteomic approach, which gave further insight into the tissue-specific ribosome RP signature. Finally, we investigated the origin of RP variations in ribosome fraction of the different tissues, by comparing RP relative abundances in our proteomic dataset and in three independent transcriptomic datasets. Interestingly, we found that, in some tissues, the RP abundance in purified ribosomes does not always correlate with the corresponding RP transcript level, arguing for a translational regulation of RP expression, and/or a regulated incorporation of RPs into ribosomes. Altogether, our data support the notion of a tissue-specific RP signature of ribosomes, which opens avenues to study how specific ribosomal composition provides an additional level of regulation to control gene expression in different tissues and organs.

## INTRODUCTION

A major challenge in cell biology is to understand how genes are expressed and regulated in space and time to control cell identity and specificity, leading to organism homeostasis. Genetic expression is defined by the unidirectional flow “DNA-mRNA-Protein”: after the DNA is transcribed in the nucleus, mRNAs are processed and exported to the cytoplasm to be translated into proteins. Tremendous amount of work has successfully focused on the regulation of the first step of the genetic flow (transcription of DNA into mRNA). The contribution of transcription factors recruited onto promoter and enhancer regions, chromatin accessibility to the transcriptional machinery (1,2) and epigenetic regulations (3) have been largely documented. Notably, mechanisms regulating gene transcription under normal and pathological conditions have been thoroughly described, as for example during regeneration of the nervous system (4,5) or in different human diseases (6,7). The rapid development and increasing depth of analysis of transcriptomic studies, thanks to next-generation sequencing, accelerate the understanding of the different levels of regulation of mRNA expression and processing.

In many studies, protein expression levels have been extrapolated from mRNA amounts (8,9). Surprisingly, combined analyses of transcriptomes and proteomes in various physio-pathological conditions revealed that mRNA and protein levels generally show only a very partial correlation (10–14). These results shed light on the critical role of post-transcriptional and translational regulation in the control of cell identity and function. Notably, regulation of the translational process is at play to control developmental programs, and alterations in its key regulators can have pathological consequences (15,16). Moreover, the translation of alternative proteins deriving from the same mRNA sequence (undetectable by RNA analysis) reinforces the notion that translational control is a major player in the control of gene expression (17,18).

Protein synthesis is a fundamental, high energy-consuming process in cellular life. The ribosome is the main functional unit in this reaction, decoding the information carried by mRNAs to assemble individual amino acids into proteins. It is a large complex composed of ribosomal RNAs (rRNAs) and ribosomal proteins. Even if its overall structure and function have been well-conserved during evolution (19), some differences are observed between organisms. For example, the eukaryotic ribosome has a higher number of ribosomal proteins (RPs) and its rRNAs are longer compared to their prokaryotic counterparts (20). The mammalian ribosome is composed of 80 RPs and 4 rRNAs, divided into two subunits: the large 60S subunit contains 46 ribosomal proteins (RPLs) and 3 rRNAs (28S, 5.8S and 5S), while the small 40S subunit is formed by 34 ribosomal proteins (RPSs) and the 18S rRNA. mRNA translation occurs through a tightly ordered sequence of three steps: initiation, elongation and termination. Each step involves specific factors (initiation, elongation and release factors - eIF, eEF and eRF) and aminoacyl-tRNAs. Moreover, the ribosome itself catalyzes peptide bound formation in the newly synthetized proteins (21).

Historically, the common dogma described the ribosome as a stable complex producing proteins from supplied mRNAs, with no regulatory role regardless of the physio-pathological conditions. Surprisingly, recent detailed studies of the translation process have revealed that the ribosome can play a more important role in the regulation of protein expression than initially thought (22,23). Indeed, it is suggested that variations in the molecular composition of this complex (rRNA, RPs, translation factors) will influence directly the quantity and/or the quality of translation.

Alterations of ribosome composition linked to specific defects in RPs have been identified in pathologies grouped under the term of “ribosomopathies”. Indeed about twenty different mutated RPs are linked to the Diamond Blackfan anemia, to malformations and to cancer predisposition (24,25). For example, somatic point mutations in individual ribosomal proteins impacts translational regulation and subsequent protein expression of oncogenic factors, leading to cancer cell proliferation (26,27). It also appears that the precise dosage of RPs contributes to maintain homeostasis and their imbalance could lead to pathologies such as the 5q-syndrome, a specific myelodysplastic syndrome caused by the loss of one copy of the gene coding for the ribosomal protein Rps14 (28). Specific RP variants were reported to impair translational fidelity (29,30). In addition, expression and phosphorylation of the ribosomal protein RPS6 conditions a phenotypic outcome in terms of central and peripheral nervous system regeneration capacity (31).

In physiological conditions, early transcriptomic studies revealed that RP paralogs could replace “canonical” counterparts in some organs. For example, the ribosomal protein L3-like (Rpl3l) is only expressed in skeletal muscle and controls myotube formation (32). Likewise, ribosomal protein L10-like (Rpl10l) is specific to the testis and critical for male meiotic transition (33). In addition, fine regulation of ribosome composition also appears to be involved in the spatial regulation of protein expression: in yeast, specific RP paralogs are required for localized translation of mRNAs, suggesting a ribosomal code (34). Specific RPs from the large or small ribosome subunits can be involved in the selection of mRNA subpools to be translated, as shown for Rpl10a (35), Rpl40 (36), or Rps25 (35). This may occur through translational regulatory elements located in transcript untranslated regions (UTR). Of example, during mammalian development, ribosomal protein Rpl38 drives specific translation of HOX genes (37–39). Moreover, the RP content of ribosomes is distinct between monosomes and polysomes (40), and the modular RP composition of ribosomes controls the translation fidelity and efficiency (41).

In addition, rRNA can also participate to translation regulation. For instance, rRNA is highly modified with more than hundreds of 2’-O methylation and pseudourydylation. Dysregulation of these rRNA modifications contributes to cancer development via alteration of translational fidelity or modification of CAP/IRES dependent translational initiation of specific mRNA (42–45). In addition, heterogeneity in the expansion segment of the rRNA could also directly control mRNA translation such as HOXa9 mRNA (46,47).

Altogether, these studies pioneered the emerging concept of “specialized ribosomes” and provide a strong link between ribosome heterogeneity, translation specificity, and specific phenotypic traits.

Despite evidences that one or even several RPs can be modified or exchanged in specific physio-pathological conditions (48), there is still a lack of comprehensive data describing ribosome heterogeneity in different cell types and tissues. Datasets comparing ribosomes composition in different tissues are mostly derived from mRNA expression levels that cannot be used to directly infer the abundances of RPs into functional ribosomes (49,50). Recently, Li and colleagues performed a proteomic analysis of 80S monosomes from nine tissues, providing evidence of variability in RP composition between the analyzed tissues (51). Alternatively, Alkan et al. developed a prediction tool based on Ribo-Seq data-extracted rRNA fragment positioning in the ribosome to highlight differential incorporation of individual RPs among 6 adult and embryonic mouse organs (52).

In our study, we purified the ribosomal fraction from 14 different adult mouse tissues and analyzed their protein composition using quantitative mass spectrometry (MS)-based proteomics. We show that ribosomes exhibit heterogeneity in RP composition, not only in the case of RP paralogs (e.g. in muscle and testis, consistent with previous works based on transcriptomics), but also for several canonical RPs displaying relative abundance variability among organs. Finally, we compared our proteomic results with transcriptomic datasets to decipher the origin of such specialization. Altogether, our work emphasizes the tissue-specific modulation of the RP content of ribosomes, and opens the way to the study of its role in the regulation of gene expression.

## RESULTS

### Characterization of ribosomal fractions prepared from different adult mouse tissues

To reveal any heterogeneity in RP composition within ribosomes across different tissues, we purified and analyzed the ribosomal fraction of different organs of wild-type (WT) adult (6 week-old) mice (**Fig 1A**). 14 different tissues dissected from 11 different organs were studied: lungs, kidneys, adrenal glands, liver, small intestine, spleen, testis, two types of muscle (heart and quadriceps skeletal muscle), various brain regions (cortex, hippocampus, olfactory bulbs), cerebellum and retina. For each tissue, we analyzed three independent biological replicates (N=3 mice, except for small organs for which we pooled several animals). After tissue lysis, ribosomes were purified by centrifugation through a sucrose cushion as previously described (53) (**Fig 1B**). Our aim is to obtain the most purified ribosomal fractions, regardless of the purity of other fractions. We confirmed by Western blot analysis that the prepared ribosomal fractions were enriched in RPs and contained no or limited contamination using several subcellular fraction markers, e.g. Histone 3 (H3) for the nuclear fraction, Hsp60 for the mitochondrial fraction, GAPDH for the cytoplasmic (post-ribosomal) fraction, and Rps6 and Rpl22 as ribosome components (**S1A Fig**). The efficiency of our ribosome enrichments was also verified by Coomassie blue staining of proteins upon SDS-PAGE separation. For each organ, the profile of the ribosomal fraction was distinct from that of the total fraction and exhibited a strong enrichment in low- to medium-molecular weight proteins that correspond to RPs (11kDa to 47kDa in mammals (54)) (**S1B Fig**).

**Fig 1:**
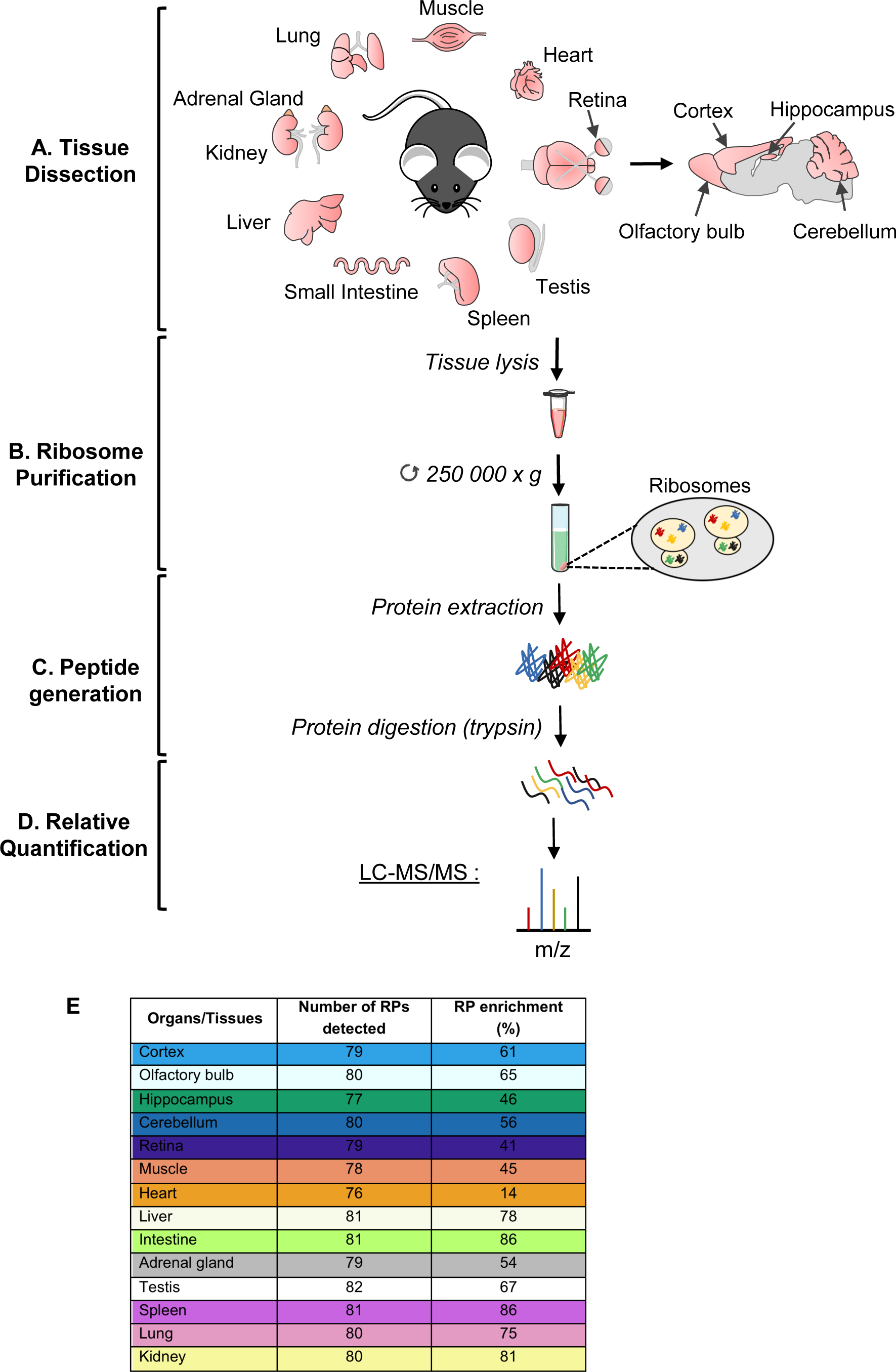
Workflow of MS-based proteomic analysis of ribosomal fractions purified from adult mouse organs. (A) Different wild-type adult mouse tissues were dissected out, lysed and processed for fractionation. Each tissue sample from one mouse corresponds to one individual biological replicate. Created with BioRender.com. (B) After separation of the nuclear and mitochondrial fractions, the ribosome fraction was purified on a sucrose cushion with ultracentrifugation at 250,000 x g for 2h. (C) Proteins were extracted from the ribosome fraction, loaded on a Bis-Tris polyacrylamide gel and in-gel digested using trypsin. (D) Extracted peptides were analyzed by liquid chromatography coupled to tandem MS for identification and relative quantification of proteins. (E) Summary of the number of RPs reliably detected in at least two replicates (RP) and the percentage of RP enrichment (Enrichment) for each tissue.

To further characterize the ribosomal fractions prepared from the different organs and tissues, their protein content was analyzed by nano liquid-chromatography (LC) coupled to MS-based quantitative proteomics (**Fig 1C-D**). 85 different RPs, 36 RPSs and 49 RPLs, were identified in at least three replicates of one tissue (**S1 Table**). For each tissue, the number of RPs reliably detected in at least two replicates ranged from 76 to 82 (**Fig 1E**). Among them, several paralogs were identified: Rpl10l, Rpl22l1, Rpl39l, Rpl3l, Rpl7l1, Rps27l and Rps4l (**S1 Table**). As calculated using the intensity-based absolute quantification (iBAQ) metrics (10), RPs accounted for more than 40% of the total protein amounts in ribosomal fractions prepared from most tissues, and notably >80% for intestine, spleen and kidney, indicating a strong enrichment in ribosomes. Only the heart ribosomal fraction presents a lower RP enrichment, reaching only 14% in our different purification attempts (**Fig 1E**).

To assess the reproducibility of our workflow, we analyzed the correlation of the measured abundances of the different RPs in the different replicates prepared from each tissue (**Fig 2A**, **S2 Fig**). The correlation coefficients indicate a high consistency in the RP abundances measured in biological replicates for all analyzed tissues. These results further confirmed the efficiency of our purification procedure to reproducibly enrich ribosomes from the different organs and tissues.

**Fig 2:**
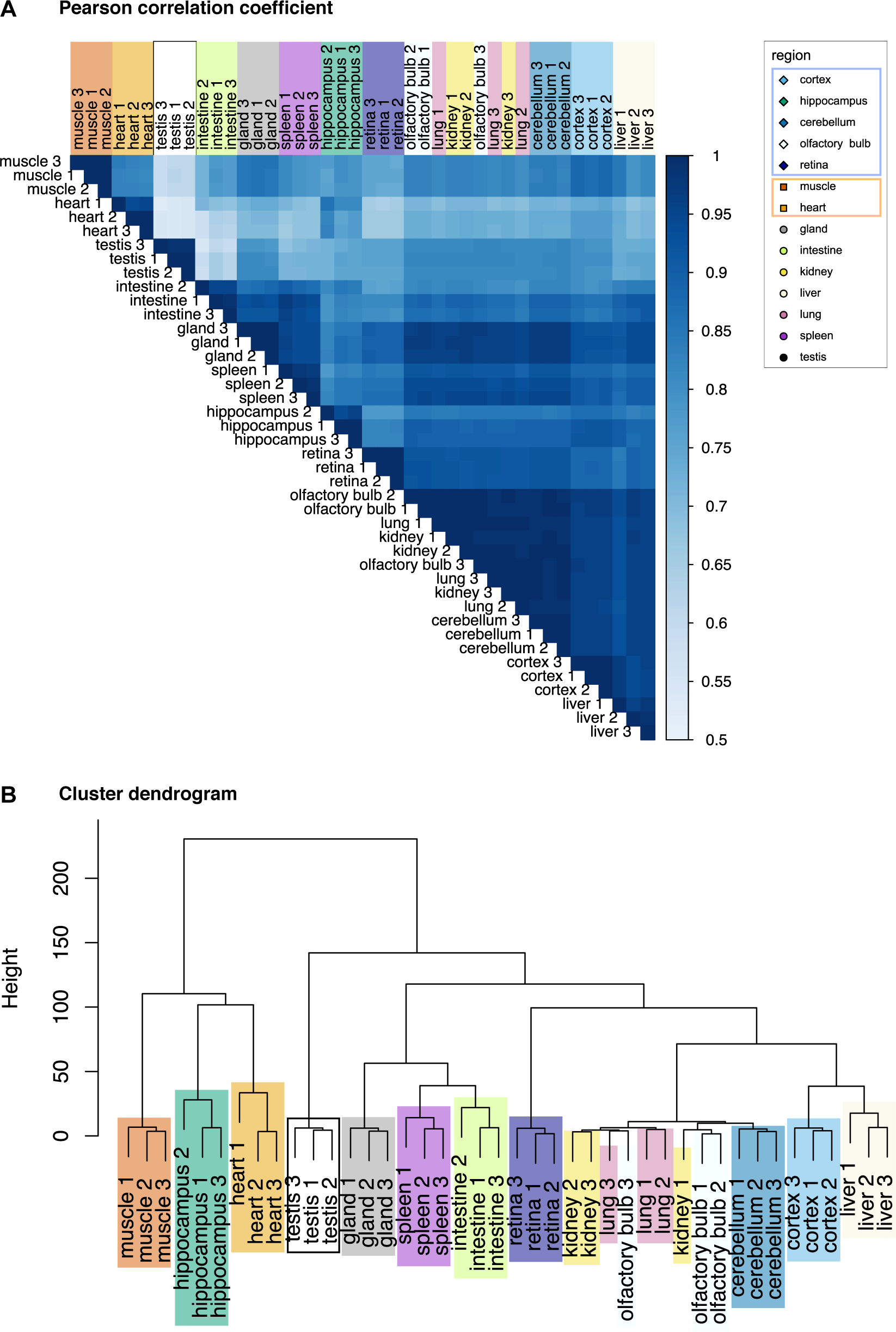
Clustering of ribosomal fractions from adult mouse organs based on RP normalized abundances. (A) Matrix of correlation showing clustering of biological replicates of ribosomal fractions from the 14 adult mouse tissues (Pearson correlation coefficient). (B) Dendrogram showing hierarchical clustering of biological replicates of ribosomal fractions of the 14 adult mouse tissues. Hierarchical clustering is computed from the euclidian distance between the log-transformed relative abundances of RP, with the Ward’s clustering method.

### RP composition as a signature of ribosomal fractions specific to adult organs and tissues

We then studied the distribution of individual RPs in the ribosomal fractions enriched from the different tissues. For each RP, we normalized the extracted abundances to the mean abundance of all samples and performed a clustering analysis after log-transformation. The dendrogram representation revealed that the three biological replicates of each tissue generally clustered together better than with the replicates of the other tissues (**Fig 2B**). This suggests some heterogeneity in the relative abundance of RPs in ribosomal fractions from different tissues.

This is exemplified by the absence of detection of a few canonical RPs in some tissues, as Rps15 detected in all tissues but the adrenal gland, intestine and muscle, and Rps23 detected in all but hippocampus and heart (**S1 Table**). This differential detection among organs/tissues was exacerbated for several paralogous RPs that were only detected in a limited number of tissues, e.g. Rpl3l specifically detected in heart and muscle, and Rpl39l and Rpl10l specifically detected in testis.

Heterogeneity in RPs distribution among the ribosomal fractions of different tissues was visualized by representing the relative abundance of each RP in the different tissues normalized by the sum of abundances of all RPs (**Fig 3**). Hierarchical clustering revealed that the majority of RPs are invariable among tissues, most of them being canonical RPs belonging to both the large and small ribosomal subunits (e.g. Rps2, Rps6, Rpl22, Rpl10a). In contrast, we also highlighted variable RPs across tissues. These variable RPs can be divided into three groups: one group containing most paralogs of canonical RPs (e.g. Rpl3l, Rpl10l, Rpl39l) that were specifically detected in one or two tissues, one group with high variability across most tissues (e.g. Rps15, Rplp1, Rpl39), and one group with variability in specific tissues (e.g. Rps26, Rps29, Rplp2) (**Fig 3**).

**Fig 3:**
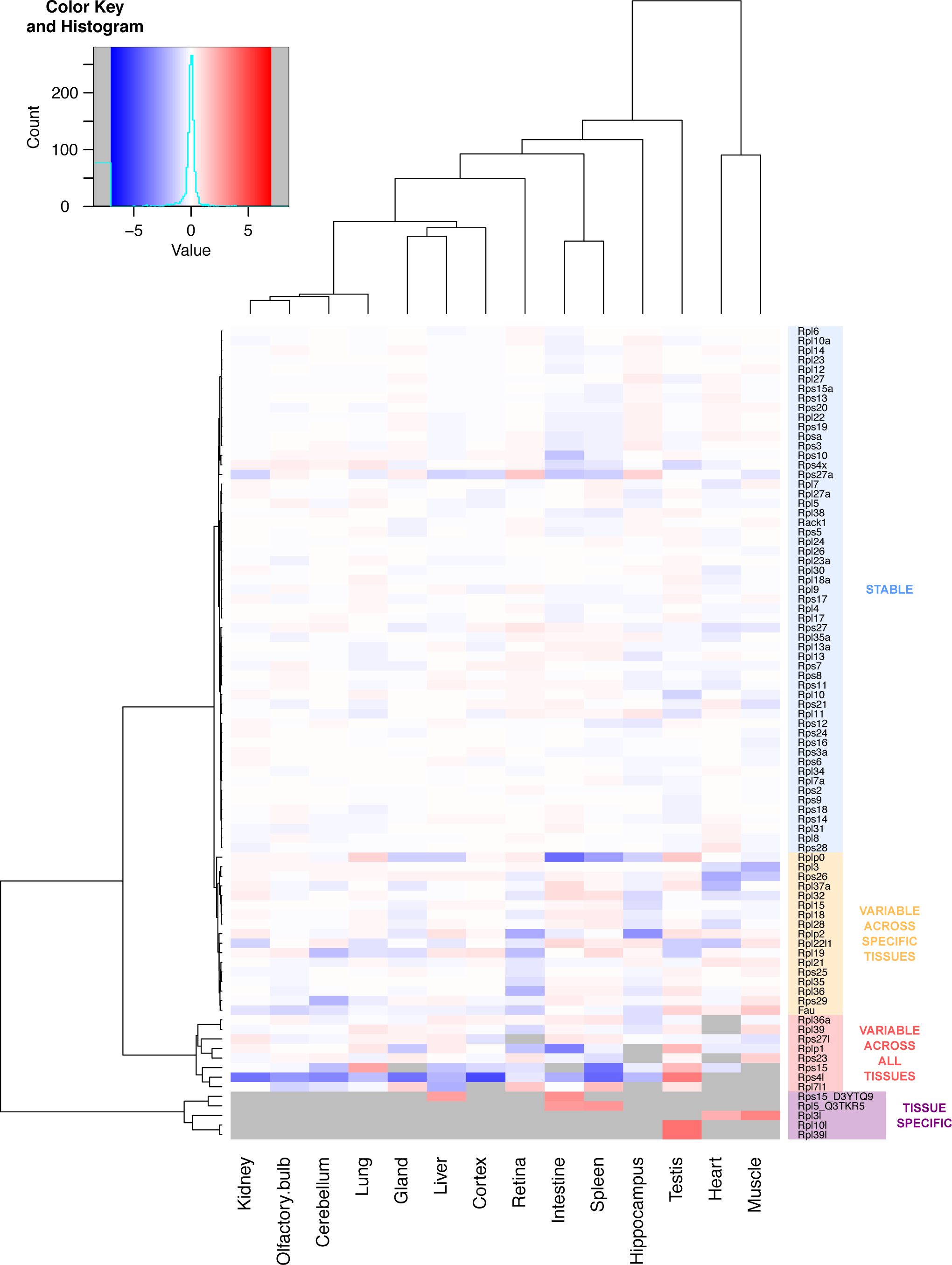
Differential RP composition of ribosomal fraction in adult mouse tissues. Heatmap of the log-transformed relative abundance normalized to the sum of all RPs for the 85 RPs detected in the ribosomal fraction of all tissues, from depleted (in blue) to enriched (in red). Grey boxes represent no detection of the RP in the ribosomal fraction of the corresponding tissue.

This heterogeneity in RP composition between ribosomal fractions of different tissues was confirmed after statistical analysis of the data. For this purpose, we used an ANOVA test, followed by One versus All LIMMA test (cf **Materials and Methods**) for RPs exhibiting ANOVA test q-value < 0.01, allowing us to highlight RPs exhibiting a significant enrichment or depletion in the ribosomal fraction of specific tissues. This strategy revealed that 59 proteins have a similar abundance in the ribosomal fractions from the different tissues analyzed (ANOVA test q-value > 0.01, or ANOVA test q-value < 0.01 but LIMMA test q-value > 0.01 or abs(log_2_FC) < 1.0) (**S2-S3 Tables**, **Fig 4A**). Among them, 46 were also highlighted in the clustering analysis as stable (**Fig 3**, group “Stable”). On the other hand, 26 RPs displayed significant variability across the different tested tissues (ANOVA test q-value < 0.01, LIMMA test q-value < 0.01 and abs(log_2_FC) > 1.0) or tissue specificity (ANOVA test q-value < 0.01, specifically detected in one or two tissues) (**Fig 4B**, **S3 Fig**, **S3 Table**). Among the specific group, 22 RPs were also highlighted in the clustering analysis as variable (**Fig 4C**, **S3 Fig**). Results from the hierarchical clustering and the statistical test were similar, strengthening the identification of heterogeneity in RP composition of ribosomes in the different organs and tissues. In the group of variable RPs, we found RPs whose normalized abundances are highly variable across all tissues, e.g. Rps30 (Fau) enriched in muscle and testis and depleted in retina, and Rplp2 enriched in liver and testis and depleted in hippocampus and retina (**Fig 4B**). We also found RPs whose normalized abundances are significantly different in a subset of specific tissues, e.g. Rps10 less abundant in intestine, Rps26 less abundant in muscle and heart, Rps29 less abundant in cerebellum, and Rpl36 less abundant in retina (**Fig 4B**, **S3 Table**).

**Fig 4:**
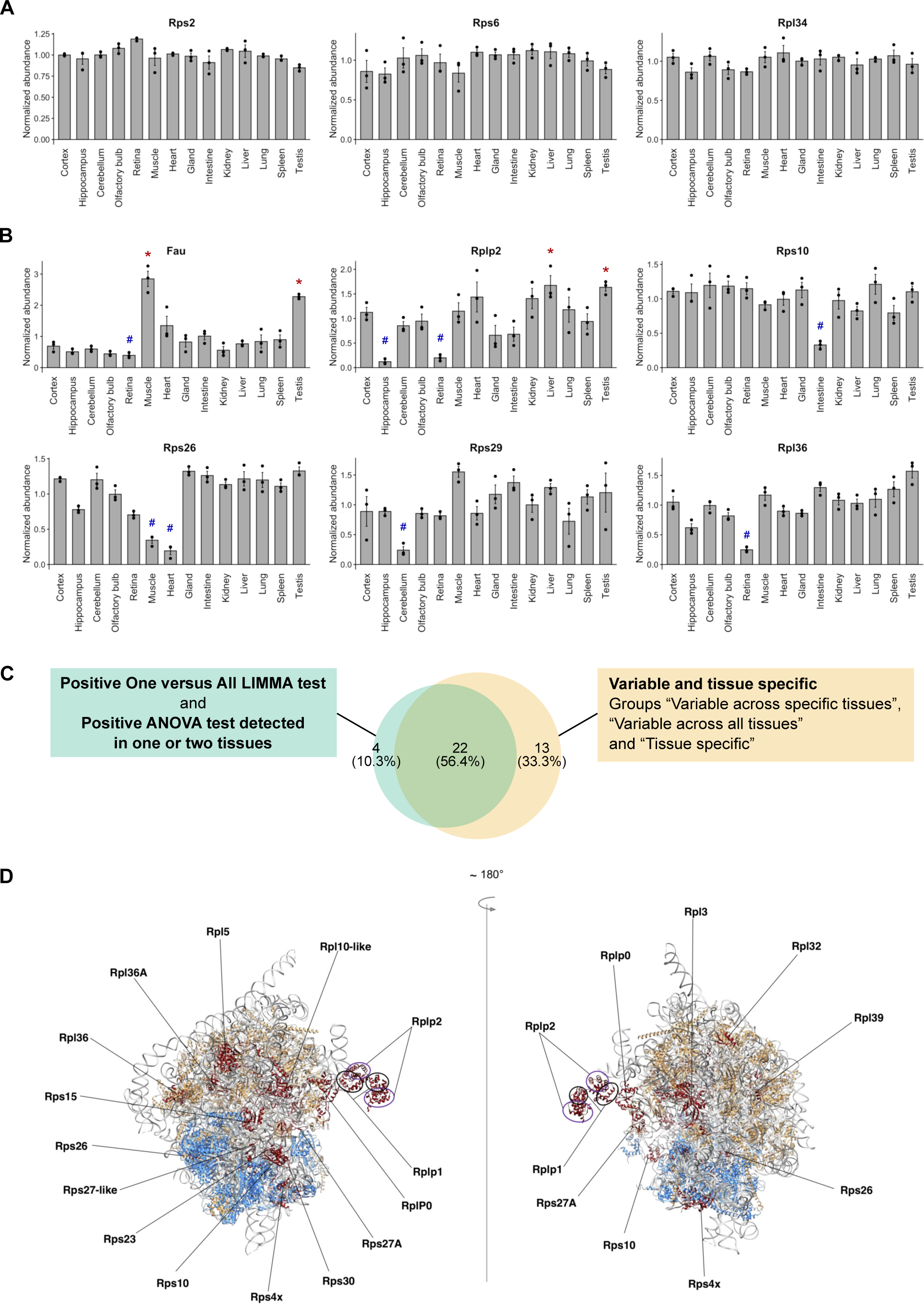
MS-based label-free quantitative proteomics shows differential abundance of individual RPs in ribosomal fractions of adult mouse tissues. (A) Barplot representation of the relative abundance in each tissue of stable RPs normalized to the sum of all RPs (Rps2, Rps6 and Rpl34 serve as examples). The mean +/- s.e.m. is plotted for each tissue, as well as values of individual replicates. (B) Barplot representation of the relative abundance in each tissue of variable RPs normalized to the sum of all RPs (Fau, Rplp2, Rps10, Rps26, Rps29 and Rpl36 serve as examples). The mean +/- s.e.m. is plotted for each organ, as well as values of individual replicates. * q-value < 0.01 (LIMMA test) and log_2_FC > 1 for One versus All comparisons. ^#^ q-value < 0.01 (LIMMA test) and log_2_FC < -1 for One versus All comparisons. (C) Venn diagram showing overlap of RPs found variable in relative abundance between tissues using statistical tests (green) and hierarchical clustering (yellow). (D) Visualization of molecular structure of the ribosome; positions of variable RPs are indicated.

Several paralog RPs have been shown to display tissue-specific transcript expression (49), as well as to control cell type-specific function, e.g. Rpl10l in the testis (33). Here, the results of our MS-based quantitative proteomic analysis demonstrated the tissue-specific protein expression and incorporation into ribosomes of these paralogs. Indeed, Rpl3l is uniquely detected in the ribosomal fractions of heart and muscle, while Rpl10l is uniquely detected in the testis ribosomes (**S4A Fig**). Interestingly, we found that the corresponding canonical RP shows decreased abundance specifically in the tissue in which the paralog version is detected. This result strongly suggest that the paralog replaces the canonical RP within the cytoplasmic ribosome. For example, Rpl3 is 2-fold and 4-fold less abundant in heart and muscle ribosomes, respectively, compared to all other tissues; and Rpl10 is 2-fold less abundant in testis ribosomes than in other tissues (**S4A Fig**).

### Position of variable RPs within the ribosome

Using Chimera software and PDB database (4v6x Human ribosome), we analyzed the localization of RPs showing variable association with the ribosome in the different tissues within the quaternary structure of the complex. Variable RPs appeared to be present either at the periphery of the quaternary structure of the ribosome (solvent side, e.g. Rps15) or at critical functional sites such as the mRNA entry site (e.g. Fau, Rps10), the tRNA binding sites: aminoacyl (A)-site (e.g. Rpl3/Rpl3l), the peptidyl (P)-site (e.g. Rpl10/Rpl10l) or the exit (E)-site (e.g. Rps26), the nascent polypeptide exit tunnel (e.g. Rpl39) and the ribosomal stalk (Rplp0, Rplp1, Rplp2) (**Fig 4D**).

Paralog versions of RPs substitute for the corresponding canonical versions in the quaternary structure of ribosomes, where it may sustain a cell type-specific control of translation. In mouse, the paralogous Rpl10l and the canonical Rpl10 differ by 3 amino acids in their sequences (**S4B Fig**), producing nearly identical 3D conformations (**S4C Fig**). Interestingly, we observed that Rpl10l occupies the exact same place as Rpl10 in the quaternary structure of the ribosome (**S4D Fig**). This suggests that a single ribosome cannot contain Rpl10 and Rpl10l at the same time, pointing towards a specialization of the ribosome through specific RP composition and suggesting downstream control of translation in different cell types and tissues. In drosophila, such paralog-switching events are a hallmark of adult gonads (55), with unique, non-overlapping functions of the paralogous and the canonical versions, as it was shown for Rpl22 and Rpl22-like (56).

### Targeted proteomics-based quantification of RPs in adult tissues

To further confirm the specific RP signature of adult organs highlighted in the label-free, unbiased proteomic approach, we performed a targeted profiling of selected RPs to analyze the ribosomal fractions from different tissues. We selected proteins from the stable group (Rps2 and Rpl34), paralogous/canonical RPs with balanced enrichment in specific tissues (Rpl3/Rpl3l, Rpl10/Rpl10l, Rpl39/Rpl39l pairs) and 4 RPs from the 22 proteins highlighted as variable from both the clustering and the statistical approaches (Rps26, Fau, Rpl36 and Rplp2) (**Fig 4B**). We deployed an isotope dilution strategy, using heavy isotope-labelled peptides whose sequences were designed based on the detection of RP-specific peptides in our label-free approach (**Fig 5A**). In total, 29 peptides were targeted in the different tissues (**S4 Table).** To be able to compare the abundances of the different targeted peptides and proteins across the different samples, we used the Rps2-derived peptide GTGIVSAPVPK as normalization reference since Rps2 was found to be stable in the ribosomal fractions across different analyzed tissues (**Fig 4A**). Of note, four selected peptides gave noisy data for which no conclusion could be drawn. Additionally, for some RPs, we observed that the absolute quantity in amol inferred from the ratio between heavy and endogenous peptide signals could vary substantially among their different peptides (**S4 Table**). This is most probably due to uneven solubilization of heavy isotope-labelled peptides, variability in digestion efficiency or limited accuracy of the heavy peptide dosage provided by the manufacturer. Nonetheless, when averaged across all selected organs, we found a high consistency among relative amounts of the measured proteins, as detailed below.

**Fig 5:**
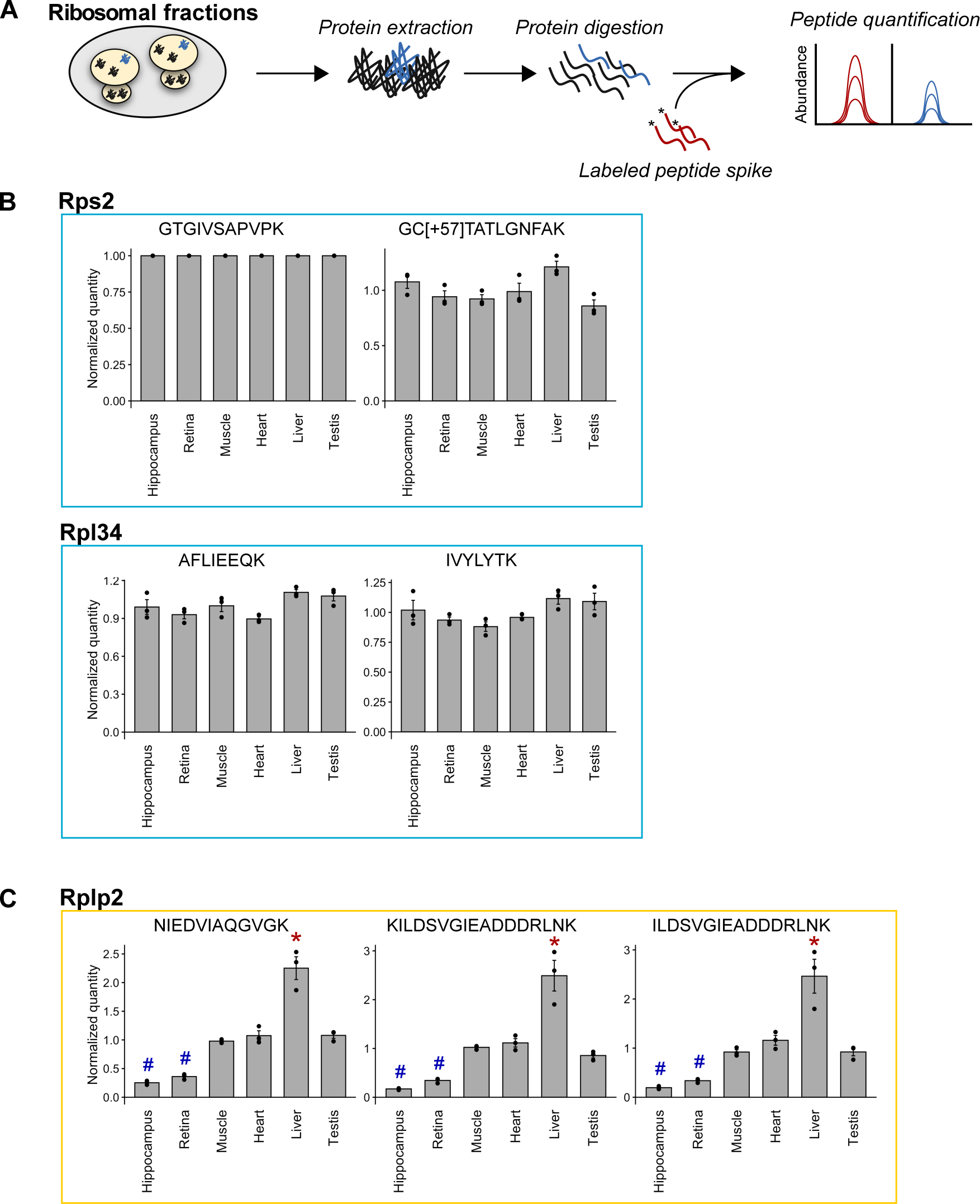
Validation of variable RPs using targeted proteomics. (A) Workflow of targeted proteomic quantification of peptides from specific proteins in ribosomal fractions of selected tissues. (B) Barplot representation of the relative abundance of stable RP-specific peptides (Rps2, taken as a reference, and Rpl34). (C) Barplot representation of the relative abundance of Rplp2-specific peptides. The mean +/- s.e.m. is plotted for each organ, as well as values of individual replicates. * q-value < 0.01 (LIMMA test) and log_2_FC > 1 for One versus All comparisons. ^#^ q-value < 0.01 (LIMMA test) and log_2_FC < -1 for One versus All comparisons.

We focused our statistical analysis on a subset of tissues (hippocampus, retina, muscle, heart, liver and testis) for which the set of RPs analyzed by targeted proteomics showed significant variations in the label-free approach. Using the targeted approach, five peptides corresponding to Rpl3l, Rpl10l or Rpl39l were specifically detected in only one or two tissues. The 19 other peptides were quantified in the 6 tissues. To decipher any significant difference in abundance between the different tissues, we performed an ANOVA test on the measured amounts of these 19 peptides. From this, 9 peptides showed no significant change in abundance between different tissues (ANOVA test q-value > 0.01, **S5 Table**). As expected, the peptides belonging to Rps2 and Rpl34 showed no significant change between tissues (ANOVA test q-value > 0.01, **Fig 5B**, **S5 Table**), confirming the stable embedding of Rps2 and Rpl34 in ribosomes from all tissues, as observed in the label-free approach (**Fig 4A**). For the 10 peptides showing significant changes between tissues (ANOVA test q-value < 0.01, **S5 Table**), we further conducted a One versus All LIMMA test to highlight which tissues had a significant enrichment or depletion of each of the corresponding proteins in their ribosomal fraction. For 9 of them, a significant depletion or enrichment was shown (comparison tissue versus all others LIMMA test q-value < 0.01 and abs(log_2_FC) > 1.0, **S5 Table**). They belong to four different proteins: Rpl10, Rpl3, Rpl36 and Rplp2. Interestingly, among them, Rpl3 and Rpl10 have tissue-specific paralogs, respectively Rpl3l detected only in the ribosomal fraction of muscular tissues, and Rpl10l detected only in the ribosomal fraction of testis. Importantly, the canonical forms were found significantly depleted in tissues in which the paralogs were specifically detected (**S5 Table, S5A-B Fig**). Measurements performed on peptides common to the paralogous and canonical forms showed a stable abundance across all organs (ANOVA test q-value > 0.01, **S5 Table, S5A-B Fig**). These results confirmed the specific RP signature of the ribosomal fractions in the muscular tissues and the testis, in which Rpl3l/Rpl3 and Rpl10l/Rpl10, respectively, balance each other and act as markers of the RP composition in the corresponding tissues.

Finally, our targeted analysis confirmed the variations in abundance of Rplp2 previously observed in ribosomal fractions from different tissues. Indeed, using three different peptides, a significant depletion of Rplp2 was detected in the ribosomal fractions prepared from the hippocampus and the retina, compared to the other organs (ANOVA test q-value < 0.01, comparison one tissue versus all others LIMMA test q-value < 0.01 and log_2_FC < - 1.0), and a significant enrichment in the ribosomal fraction of the liver (ANOVA test q-value < 0.01, comparison one tissue versus all others LIMMA test q-value < 0.01 and log_2_FC > 1.0) (**Fig 5C**). These results were consistent with those of the label-free approach and further confirmed the differential RP composition of ribosomal fractions in the different tissues.

### Correlation of relative transcript expressions and RP composition of ribosomes in adult tissues

To unravel the origin of observed variations in RP content of ribosomal fractions from different adult mouse tissues, we sought to analyze RP expression at the mRNA level in each organ. For this, we used published datasets mouse transcriptome atlas: the transcriptomic BodyMap (57), the Mouse ENCODE Consortium project (58)). We also use an Human transcriptome atlas (the Illumina Human Body Map (GSE30611)) in order to see to which extend our results in mouse can be extrapolated in human. We compared the level of RP expression from our proteomic analysis to each of these 3 mRNA expression dataset as analyzed by (49). To estimate RP transcript abundances, we computed the reads per kilobase per million mapped reads (RPKM) normalized by the sum of RPKM of all the RPs in each tissue (49). The relative expression of RPs among organs represented as a heatmap shows little variation at the transcript level, except for the three paralogous RPs: Rpl3l, strongly enriched in the heart and muscle, and Rpl10l and Rpl39l, strongly enriched in the testis (**S6 Fig**).

We then compared the expression of each RP in the three transcriptomic datasets (“Mouse BodyMap”, “Mouse ENCODE” and “Human Body Map”) with the results of our MS-based quantitative proteomic analysis of ribosomal fractions. We focused on organs and tissues found in all four datasets: brain, gland, heart, kidney, liver, lung and testis. For the Mouse ENCODE dataset, no direct data from cortex are available as they generated a brain sample. We decided then to compare values of the brain from that of the cortex.

For many of the stable RPs found in our proteomic analysis (59 RPs with non-significant difference in abundance in ribosomal fractions of the different tissues), the corresponding transcript expression did not vary across organs in all transcriptomic datasets, e.g. Rps12, Rps16, Rpl14 and Rpl34 (**Fig 6A**), showing an overall good correlation between transcriptomic and proteomic levels for this group of RPs. Moreover, several variable RPs detected by our proteomic approach (26 RPs with significant variation in abundance in ribosomal fractions of the different tissues) also showed variability at the transcript level. This is the case of the paralogous RPs Rpl3l, Rpl10l (not reported in the Mouse ENCODE dataset), Rps4l (not reported in the Mouse ENCODE and the Human Body Map datasets) and the corresponding canonical Rpl3, Rpl10 and Rps4x (**Fig 6B**). Interestingly, the relative transcript expression of the canonical RPs corresponding to Rpl3l and Rpl10l correlates with the relative abundance measured in our proteomic data. Indeed, Rpl3 transcript is less abundant in the heart than in the other organs, while Rpl10 is less abundant in the testis (**Fig 6B**).

**Fig 6:**
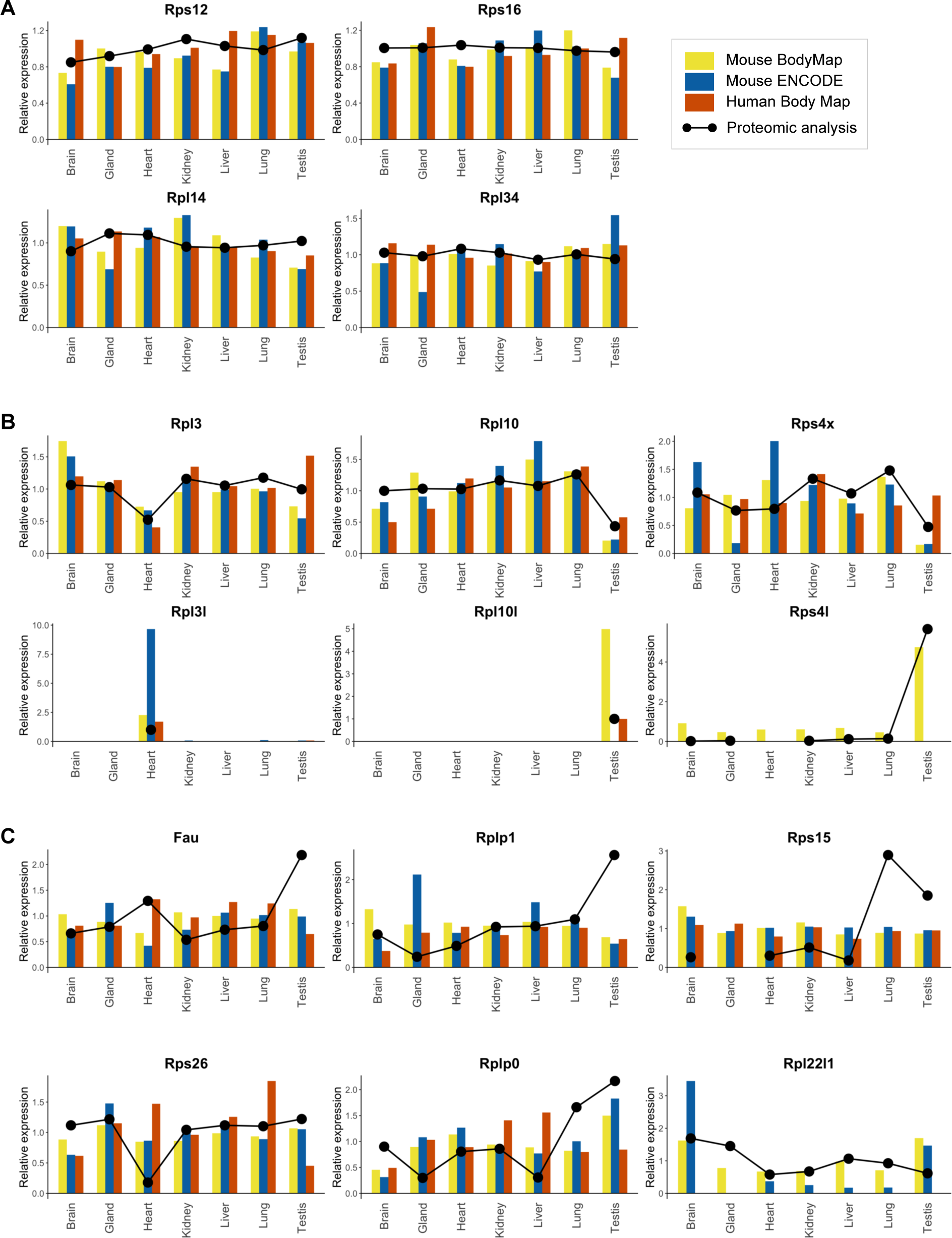
Comparison of the relative expression of RPs in transcriptomic-based datasets and MS-based proteomic analysis of the ribosomal fraction in different mouse tissues. (A-C) Barplot representation of the relative expression of each RP at the transcript level, with data from Mouse BodyMap (57), Mouse ENCODE Consortium (58) and Illumina Human Body Map 2.0 as analyzed by (49)). The relative abundance as determined in our present study is superimposed with black dots and line. (A) Examples of stable RPs with similar transcriptomic and ribosome proteomic profiles (Rps12, Rps16, Rpl14, Rpl34). (B) Examples of paralogous RPs and corresponding canonical forms, with similar transcriptomic and ribosome proteomic profiles (Rpl3l/Rpl3, Rpl10l/Rpl10, and Rps4l/Rps4x). (C) Examples of variable RPs with tissue-specific enrichment or depletion in the ribosomal fraction not reflected by the transcriptomic profile (Fau, Rplp1, Rps15, Rps26, Rplp0, Rpl22l1).

In contrast, for several variable RPs, we observed little correlation between the relative expression level of transcripts in tissues and the relative abundance of corresponding proteins in the ribosomal fractions. This is the case for Fau and Rplp1, both found enriched in the ribosomal fraction of the testis at the protein level, but measured at similar amounts in the different tissues at the transcript level (**Fig 6C**). Similarly, Rps15 was found significantly enriched in ribosomal fractions of the lung and testis compared to other tissues, but its transcript level was found stable across tissues. Another example is Rps26 found in lower protein amounts in the ribosomal fraction of the heart compared to other tested organs/tissues, but found in similar amounts at the transcript level (**Fig 6C**). In other cases, transcript levels showed consistent enrichment or depletion in a particular tissue, but with a decorrelation from the abundances measured in ribosomal fractions for the corresponding proteins. This is the case of Rplp0, found enriched in the ribosomal fractions of the lung and the testis, but whose transcript level is consistently depleted in the brain (**Fig 6C**). Another example is Rpl22l1, depleted in heart and testis ribosomes, but enriched in the brain and testis at mRNA level.

Altogether, these results indicate that the relative level of transcripts of a given RP does not necessarily correlate with the relative abundance of its corresponding protein in the ribosomal fraction in adult organs/tissues. Even if these conflicting results may be due to the analysis of different biological samples, they bring out interesting hypotheses: (i) differential post-transcriptional regulation of RP protein expression among organs, and/or (ii) differential incorporation of specific RPs into ribosomes among organs. This may lead to an organ-specific control of translation, which remains to be firmly demonstrated.

## DISCUSSION

The ribosome was generally regarded as an invariable machinery producing proteins from mRNA templates. It was considered as not being directly involved in the regulation of translation or the selection of mRNAs to be translated. Yet, this dogma has been challenged over the past few years, notably by an accumulation of compelling evidence supporting the variability of ribosome composition across different cell types and in various physiological and pathological conditions (59–61). Here, we used MS-based quantitative proteomics to decipher the differential RP composition of ribosomes in 14 different tissues across 11 organs of adult mouse. While the majority of RPs showed no significant difference in abundance in ribosomal fractions prepared from different organs, we found several RPs with variable levels. Indeed, some RPs are clearly enriched or, on the other hand, depleted from specific tissues or organs (**Fig 3**), which supports the concept of specific ribosomal signatures of adult mouse organs (**Fig 7**). This is especially striking given that our macroscopic approach analyzed ribosomal fractions of whole-tissue or whole-organ, strongly suggesting that variations in RP composition likely control organ- or tissue-specific development and function. Similar approach at a whole-organ level in drosophila highlighted specific RP clusters in the 80S fraction of ovary and testis, mainly driven by RP paralog enrichment or switching in the ribosome (55).

**Fig 7:**
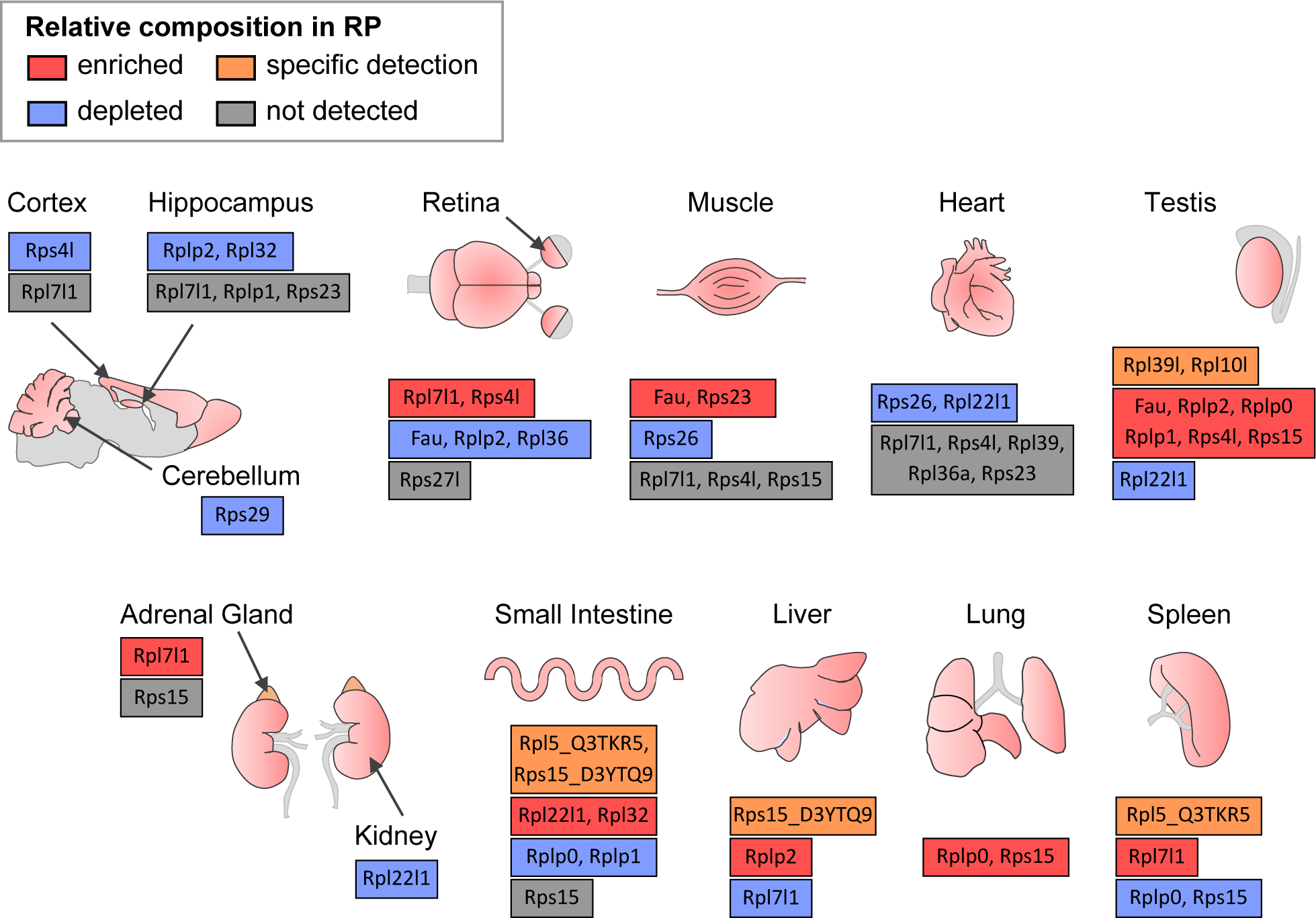
Summary of variations detected for individual RPs in the ribosomal fraction of different mouse tissues and organs. Relative RP enrichment and depletion among the mouse tissues and organs, as well as specific detection and absence of detection, are illustrated for the 22 RPs highlighted as variable from both the clustering and the statistical analyses (see **Fig 4C**).

We observed that numerous variable RPs identified by our work are located in the peripheral regions of the ribosome. So, we cannot rule out the possibility that these RPs may be detached from the ribosome during sample processing, depending on the tissue. Yet, as we treated all tissues and organs in the same way, we should expect similar behaviours, unless the strength of the interaction of some RPs with the ribosome is different from tissue to tissue. Therefore, we account for their differential detection not by a possible detachment of these RPs during sample preparation, but rather by their actual differential integration into the ribosomal complex. With this assumption, we detected significant differences in ribosome composition among the different organs, suggesting an organ-specific RP signature of the ribosome.

With the ambition of determining the absolute amounts of specific RPs within ribosomes, we used a targeted proteomics approach relying on the use of isotopically-labelled peptides. Unexpectedly, we found a major limitation: the peptides quantities determined could differ from a factor of 10 between different measured peptides of the same RP. Even if this issue prevented us from concluding on the precise stoichiometry of each RP within the ribosomal fraction of each tissue, we found great consistency between relative amounts of the different peptides of each RP in the different tissues. These results further confirm the significant differences in RP composition highlighted by the label-free approach. It is interesting to note that other studies using a similar targeted strategy have based their conclusions on the analysis of at least two peptides per protein, or of a single peptide with an additional control for specificity of detection (35).

Our proteomics data on various adult mouse tissues support the concept of heterogeneity in ribosome composition, which may underlie the specific regulation of translation. Considering the variability in the RP composition of the ribosomal fraction in the different tissues analyzed, our observations are suggestive of a specialized function of variable (tissue-enriched or tissue-depleted) RPs in the regulation of the translational process to serve specific functions. This feature may be linked to specialized translation that ensures cell type-specific functions, which remains to be demonstrated for many RPs. This is particularly the case of Rpl39l, a paralog of Rpl39, consistently found to be enriched in the testis. A recent study used a proteomics-based approach, similar to ours, to highlight inter-organ variations in 80S monosome composition, and showed Rpl39l enrichment in sample prepared from the testis (51). Even though this analysis yields to different clustering of variable versus stable RPs compared to our results, potentially due to the analysis of different fractions (*i.e.* 80S monosomes and total ribosomes), this study emphasizes that the differential integration of Rpl39l in ribosomes, not only in adult organs, but also over the course of development, has a direct impact on male germ cell function (51).

At the cellular level, ribosome heterogeneity may allow cells to respond rapidly to external stimuli by regulating gene expression at the translational level (22). Whether the intracellular population of ribosome is itself heterogeneous is an important yet technically challenging question to address. It has been tackled by Barna and colleagues in mouse embryonic stem cells (35). By using MS-based quantitative proteomic approaches to characterize translationally active ribosomes, they found RPs with variable abundances between ribosomes from polysomes and free subunits (e.g. Rpl10a/uL1, Rpl38/eL38, Rps25/eS25), as opposed to invariant RPs. These results support the notion of an heterogeneous population of ribosomes within the cell to sustain various cellular functions via mRNA-specific translation (35). Furthermore, advanced cryo-electron tomography imaging techniques will enable to resolve ribosome heterogeneity at a subcellular level, thus identifying intra-cellular populations of specialized ribosomes, for instance in association with different organelles (62).

Then the next key question is: what mechanisms are at play to control for inter-organ variation in RP ribosomal composition? In the present study, we compared our proteomic data to transcriptomic data from available mouse and human wide RNA-sequencing datasets. The transcript and protein relative expressions correlate well for many RPs among the different organs. This is particularly the case of paralogous RPs Rpl3l, Rpl10l and Rps4l, and the respective canonical RPs Rpl3, Rpl10 and Rps4x that, in addition, show organ-specific compensations at the transcript level as well as at the protein level. Interestingly, this phenomenon of regulation has already been demonstrated for the pair Rpl22/Rpl22l1, where Rpl22 itself has been shown to repress the translation of Rpl22l1 by binding to an hairpin structure of its mRNA (63). The finding that the paralogous version of a RP competes with the canonical version to incorporate into the ribosome reinforces the hypothesis of specific regulation of the ribosome composition. Several studies describe that such a “paralogous RP signature” ensures specific functions through selective translation, e.g. muscle function (32,64) or gonad specification and function (55,65). In yeast, the balance between two RP forms controls proper mitochondrial function (66) and response to stress (67).

An important point is to define how the expression of individual RPs is specifically controlled and what the upstream regulatory mechanisms are. Regulation of expression of individual RPs may occur at the transcription level through the control of cell type-specific transcription factors. Indeed, studying RP gene expression in different human tissues, Guimaraes and colleagues identified transcription regulatory elements located in the promoter of RP genes, suggesting that lineage-specific regulations may be at work through the activity of particular transcription factors (50). Although little difference is observed in promoter utilization in cell type-specific versus non-specific RPs, and despite the heterogeneity in promoter regulatory sequences of individual RPs, it is possible that a set of cell type-specific transcription factors orchestrate RP gene expression (50,68). This accounts for the level of heterogeneity observed at the transcript level, where mechanisms of co-regulation of RPs expression remain to be determined (37).

In a meta-analysis of several transcriptomics-based and translatomics-based studies on humans and rodents, Panda et al. highlighted organ-, stage- and tumorigenic state-specific RP signatures (69). Interestingly, they found a high correlation between ribosome-protected footprints and mRNA levels for mRNA coding for RPs, notably in liver, brain, hippocampus and heart (69), suggesting correspondence between RP mRNA and protein levels. However, taking into account extra-ribosomal functions of RPs (70) and actual integration of proteins in the functional ribosomal complex is crucial to further illustrate ribosome heterogeneity in terms of RP composition. Our study reveals that several RPs display no or little correlation between the relative transcript expression level and the relative protein abundance among different mouse tissues (**Fig 6C**). Similarly, in drosophila, the inter-organ differences in RP composition are not simply driven by variations in RP mRNA levels (55).

This observation brings out several hypotheses, such as a differential post-transcriptional regulation of RP protein expression among tissues, and/or a differential incorporation of RPs into the ribosome. These results also pave the way for the key questions, as yet to solve, of how and where the cell fine-tunes the ribosome composition to regulate translation. Of note, in our study, Rplp0, Rplp1 and Rplp2 (RPs of the ribosomal stalk, a lateral protuberance of the ribosome that binds translation factors) were found variable in both the clustering and the statistical approaches. These RPs also show inter-tissue variability at the mRNA level (71). Interestingly, free Rplp1 and Rplp2 integrate into the ribosome in the cytoplasm, and this association regulates the translational activity of the ribosome (72). More recent studies have demonstrated the local protein synthesis and integration of individual RPs in the ribosome in distal neuronal compartments (73,74), pointing towards a local remodelling of ribosome RP composition during neuronal circuit development or in response to stress.

To conclude, our study adds up to the growing evidence that diverse cell types exhibit specific – and probably specialized – RP expression in a mammalian organism. Based on MS-based quantitative proteomic data, our work brings firm evidence that this tissue-specificity occurs not only at the protein expression level, but also at ribosome integration level. Comparison with transcriptomic datasets shows that these specific RP signatures do not necessarily correlate with RP mRNA levels, opening questions about inter-tissue ribosome heterogeneity regulatory mechanisms.

## MATERIAL AND METHODS

### Tissue sampling and ribosome purification by subcellular fractionation

Wild-type (WT) C57BL/6J adult mice were used in this study, regardless of their sex, except for collection of the testis. Organs and tissues of 4 to 6 week-old mice were dissected and flash-frozen in liquid nitrogen. Ribosomal fraction purification was performed according to (53). All steps were performed on ice or at 4°C. Samples were lysed in freshly prepared buffer A (50mM Tris HCl pH 7.4, 250 mM sucrose, 250 mM KCl, 5 mM MgCl_2_ (Sigma-Aldrich)) using a Cell Mill (RETSCH MM 400). An aliquot of the cell suspension (total fraction) was saved for SDS-PAGE. IGEPAL detergent (Sigma-Aldrich) was added to the remaining volume to a final concentration of 1%. After 20 min incubation on ice, the lysate was centrifuged at 750 x *g* to pellet nuclei (nuclear fraction) then 12 500 x *g* to pellet mitochondria (mitochondrial fraction). The supernatant (post-mitochondrial fraction) was loaded on a sucrose cushion (1.25 M sucrose, 0.25 M KCl, 5 mM MgCl_2_, 50 mM Tris-HCl pH 7.4) and ultracentrifuged at 250 000 x *g* for 2 h (Beckman Optima TL 100 Ultracentrifuge). After ultracentrifugation, 50µl of supernatant (cytoplasmic fraction) was saved. The ribosome pellet was washed twice in ultrapure ice-cold water and resuspended either in 50µl of Laemmli buffer (ribosomal fraction) or in buffer C (tris HCl 50mM pH7.4; 5mM MgCl_2_; 25mM KCl). 0.5μl benzonase (Sigma-Aldrich) was added to the nuclear sample and incubated for 10 min at 37°C to digest DNA.

### Protein quantification

After denaturation at 95°C for 5 min, the protein concentration of all fractions was determined using Pierce BCA Protein Assay Kit (ThermoFisher Scientific).

### Western blot

20µg of protein were loaded on a 12% SDS-PAGE gel and separated by electrophoresis for 4h at 230V. The proteins were then transferred on nitrocellulose membrane (Thermo Scientific) under a constant amperage of 250 mA for 3 hr. The membrane was blocked in 5% milk powder in Tris-Buffered Saline (TBS) and incubated with the following primary antibody diluted in 5% milk powder in TBS with 0.1% tween (TBS-T) overnight under agitation at 4°C: anti-RPL22 (1:2000, mouse, Santa Cruz sc-373993), anti-RPS6 (1:5000, Rabbit, Cell Signaling 2217), anti-H3 (1:10 000, Rabbit, Cell Signaling 9715), anti-HSP60 (1:2000, mouse, Santa Cruz sc-376240), anti-GAPDH (1:5000, Mouse, Proteintech 60004-1-Ig). On the following day, membranes were washed then incubated with horseradish peroxidase-linked secondary antibodies: anti-rabbit (Proteintech SA00001-2) or anti-mouse (Millipore 12-349) diluted to 1:5000 or 1:10 000 in TBS-T. Membranes were probed with ECL substrate (100 mM Tris HCl pH 8.5, 0.5% coumaric acid (Sigma-Aldrich), 0.5% luminol (Sigma-Aldrich) and 0.15% H_2_O_2_ (Sigma-Aldrich)). Chemiluminescence was visualized with the ChemiDoc system (Bio-Rad).

### Coomassie staining

10 µg of protein were loaded on 12% SDS-PAGE gel and separated by electrophoresis for 4h at 230V. The gel was fixed in fixing solution (50% methanol (VWR Chemicals), 10% glacial acetic acid (VWR Chemicals)) for 1hr with gentle agitation. After fixation, the gel was incubated in staining solution (0.1% Coomassie Brilliant Blue R-250 (Bio-Rad), 50% methanol, 10% glacial acetic acid) for 20 min with gentle agitation. The gel was then washed several times with destaining solution (40% methanol, 10% glacial acetic acid). Gels were imaged when the gel’s background was fully distained.

### Discovery Proteomics

#### Mass spectrometry-based analysis

Proteins (between 5 and 10µg) from tissue preparations were solubilized in Laemmli buffer before loading on top of a 4–12% NuPAGE gel (Life Technologies), stained with R-250 Coomassie blue (Bio-Rad) and in-gel digested using modified trypsin (sequencing grade, Promega) as previously described (75). The dried extracted peptides were resuspended in 5% acetonitrile and 0.1% trifluoroacetic acid and analyzed by online nanoliquid chromatography coupled to tandem mass spectrometry (LC–MS/MS) (Ultimate 3000 RSLCnano and the Q-Exactive HF, Thermo Fisher Scientific). Peptides were sampled on a 300 μm 5mm PepMap C18 precolumn (Thermo Fisher Scientific) and separated on a 75 μm 250 mm C18 column (Reprosil-Pur 120 C18-AQ, 1.9 μm, Dr. Maisch HPLC GmbH). The nano-LC method consisted of a 60 min multi-linear gradient at a flow rate of 300 nl/min, ranging from 5 to 33% acetonitrile in 0.1% formic acid. For all tissues, the spray voltage was set at 2 kV and the heated capillary was adjusted to 270°C. Survey full-scan MS spectra (m/z = 400–1600) were acquired with a resolution of 60 000 after the accumulation of 10^6^ ions (maximum filling time 200 ms). The 20 most intense ions were fragmented by higher-energy collisional dissociation after the accumulation of 10^5^ ions (maximum filling time: 50 ms). MS and MS/MS data were acquired using the software Xcalibur (Thermo Scientific).

#### Data processing

Data were processed automatically using raw2mzDB converter version 0.9.10. Peaklists were obtained using mzDB-access version 0.8.0 (https://github.com/profiproteomics/mzdb/tree/mzdb-processing_0.8.0) by executing the ‘create_mgf’ command (parameters: MS level=2, no intensity cut-off, precursor_mz=isolation_window_extracted). Peptides and proteins were identified using Mascot (version 2.6) through concomitant searches against a home-made Mus database, classical contaminants database (homemade) and their corresponding reversed databases. The Mus database is composed from Mus musculus Reference Proteome (UP000000589 from UniProt), ribosomal proteins and their isoforms, and home-selected sequences to identify potential variants from 13 mouse canonic ribosomal protein sequences of special interest (variant obtained by blasting their mRNA sequence against the genome or their protein sequence against the translated genome). Finally, this database has been curated to be non-redundant in protein sequences. Trypsin/P was chosen as the enzyme and two missed cleavages were allowed. Precursor and fragment mass error tolerances were set, respectively, to 10 ppm and 25 mmu. Peptide modifications allowed during the search were: carbamidomethylation (fixed), acetyl (protein N-terminal, variable) and oxidation (variable). The Proline software (version 2.1) (76) was used to merge and filter results for each tissue separately: conservation of rank 1 peptide-spectrum match (PSM) with a minimal length of 7 and a minimal score of 25. PSM score filtering is then optimized to reach a False Discovery Rate (FDR) of PSM identification below 1% by employing the target decoy approach. A minimum of one specific peptide per identified protein group was set. Proline was then used to perform MS1-based label free quantification of the peptides and protein groups from the different samples without cross-assignment activated between tissue but activated only between replicates. Protein iBAQ were computed from specific peptides abundances.

Proteins were filtered out if they were not identified in the 3 replicates of at least one tissue. Protein values were discarded from a tissue if it was detected in one single replicate of a tissue. For each tissue and replicate, total ribosomal protein iBAQ was used to normalize iBAQ of quantified proteins. After log_2_ transformation of normalized iBAQ, ProStaR (77) was used to impute missing values. POV missing values were imputed with slsa method and MEC ones with a low value. For each sample, this low value was set at the minimum value observed for a protein in this sample. Statistical testing was conducted only on ribosomal proteins using ANOVA with a p-value cut-off allowing to reach a FDR inferior to 1% according to the Benjamini-Hochberg procedure. RPs quantified in one or two tissues were considered tissue-specific. For other proteins, in order to discriminate in which tissue RPs positive to ANOVA were differentially abundant, we used Prostar (77) to run LIMMA test with One versus All as contrast. All data were then merged and p-values corrected with Benjamini-Hochberg procedure. Log_2_(Fold Change (FC) One versus All) were calculated without using MEC imputed values. Proteins were considered as significantly enriched (or respectively depleted) in a tissue if their FDR-adjusted p-value was below 0.01 and their log_2_(FC) > 1 (respectively log_2_(FC) < -1). Proteins were considered as not detected in the tissue if their FDR-adjusted p-value was below 0.01 and no log_2_(FC) could be calculated.

#### Data representation and analysis

Ribosomal proteins were filtered out if they were not identified in the 3 replicates of at least one tissue. Protein value were discarded from a tissue if it was detected in one single replicate. All plots were generated using R software for data representation and analysis (78). Scatterplots of RP hits were obtained by plotting the log_10_-transformed protein abundances (iBAQ) normalized to the total iBAQ of all RPs per organ, across biological replicates of each organ. To highlight the relationship between the ribosomal fractions of the different organs, the Pearson’s correlation coefficient was used to compare samples from RP iBAQ values. To generate the heatmap of RPs, iBAQ of each RP of one organ was normalized to sum of all RP iBAQ. For each RP, the relative abundance was computed as the log_2_-transformation of the normalized iBAQ averaged on all organs. To highlight the biological differences between the ribosomal fractions of the different organs, hierarchical clustering was performed by computing the euclidian distance of the log-transformed relative abundances of RP between all samples, with the Ward’s clustering method. Barplots of the relative expression of RPs in the ribosomal fraction of each organ were obtained from the iBAQ normalized to the sum of all RPs, and averaged across all organs.

### Quaternary structures

Crystal structures from human ribosomes were downloaded from PDB (4x6v, 6oli). All structures were generated using Chimera software (79), version 1.15c. Labels were manually added.

### Targeted Proteomics

#### Mass spectrometry-based analysis

Samples were in-gel digested as described for discovery proteomics. The dried extracted peptides were resuspended in 5% acetonitrile and 0.1% trifluoroacetic acid with a mixture of heavy isotope-labeled peptides from Synpeptide (Shanghai, China) spiked-in. The sample were analyzed by online nanoliquid chromatography coupled to tandem mass spectrometry (LC–MS/MS) (Ultimate 3000 RSLCnano and the Q-Exactive HF, Thermo Fisher Scientific). Chromatographic parameters were the same as described for discovery proteomics. The targeted acquisition method combined two scan events corresponding to a full scan event and a time-scheduled PRM event targeting the precursor ions selected for the pairs of heavy and endogenous peptides in ±2.7 min elution time windows. The full scan MS spectra (m/z = 400–1600) were acquired with a resolution of 30 000 after the accumulation of 10^6^ ions (maximum filling time 200 ms). The PRM event were acquired with a resolution of 30 000 after the accumulation of 10^6^ ions (maximum filling times varying from 55ms to 260ms depending on the number of peptides to target in each run time range).

#### Data processing

Targeted data were processed with Skyline 4.2. Five best product ions (mono-charged y-type ions) per precursor ion were extracted with 5ppm tolerance. All matching scans were used. Chromatographic peaks were investigated to manually adjust peak integration boundaries transitions. Peptides whose heavy version was observed with too much variation (coefficient of variation superior to 50%) were discarded. For each tissue, only transitions detected with Signal to Noise higher than 10 and in its 3 replicates were finally used. Signal at the peptide level was obtained by summing the corresponding transition peak areas for heavy and endogenous peptide. The ratios between the endogen and the heavy peptides were used to determine the mol amount of endogenous peptides. To take into account the variability of ribosomal proteins amount in the samples, all data were normalized by the peptide GTGIVSAPVPK from Rps2. We selected tissues for which variations for the followed peptides were expected from the label-free approach (hippocampus, retina, muscle, heart, liver and testis). For peptides detected in the 6 tissues, statistical testing was conducted on log_2_ transformed data using an ANOVA with a p-value cut-off allowing to reach a FDR inferior to 1% according to the Benjamini-Hochberg procedure. To discriminate in which tissue peptides positive to ANOVA were differentially abundant, we use Prostar to run LIMMA t-test with One versus All as contrast. All results were merged and p-values corrected with Benjamini-Hochberg procedure. Peptides were considered as significantly enriched (respectively depleted) in a tissue if their FDR-adjusted p-value was below 0.01 and log_2_FC > 1 (respectively log_2_FC < -1).

#### Data representation and analysis

All plots were generated using R software for data representation and analysis (78). For each individual peptide, the relative abundance was obtained from the mol amount (inferred from the ratio between heavy and endogenous peptide signals) averaged on selected organs.

### Analysis of transcriptomic datasets

To compare the relative expression of RP at the transcript level, we used published dataset of mouse and human transcriptomic atlas of adult organs: the Mouse Transcriptomic BodyMap (57), the Mouse ENCODE Consortium project (58)) and the Illumina Human Body Map (GSE30611) as analyzed by (49) (Supplementary Table S6 of the publication). For all datasets, read per kilobase per million mapped reads (RPKM) values were retrieved. For the Mouse BodyMap, only samples from males were selected and RPKM values were averaged by organ. We selected all RPs detected in our dataset and all organs in common to the three transcriptomic datasets and our proteomic dataset, with the only apporximation of “Cortex” as “Brain” when not available. We adopted the same normalization strategy as (49) and computed the RPKM normalized to the sum of RPKM of all RP of each organ times the number of RP (RPKM * number RPs / sum(RPKM)). To generate the heatmap, we computed the log_2_-transformation of the normalized RPKM averaged on all considered organs. Barplots of the relative expression of RP transcripts of each organ were obtained from the normalized RPKM values averaged on all organs.

### Data availability statement

The discovery LC-MS/MS data have been submitted to the ProteomeXchange Consortium via the PRIDE (80) partner repository under dataset identifier PXD044060.

## ACKNOWLEDGMENTS

This work was supported by the Agence Nationale de la Recherche (ANR) grant to SB (ANR-18-CE16-0007). The proteomic experiments were partially supported by ANR under projects ProFI (Proteomics French Infrastructure, ANR-10-INBS-08) and GRAL, a program from the Chemistry Biology Health (CBH) Graduate School of University Grenoble Alpes (ANR-17-EURE-0003). JS is supported by Fondation pour la Recherche Médicale (FRM) (SPF201909009106). HN is supported by the NRJ Foundation and the European Research Council (ERC-St17-759089).

## AUTHOR CONTRIBUTIONS

SB and YC designed the project, with inputs from HN. JS, AMH, HN, SB and YC wrote the manuscript with input from all co-authors. MRB, HN and CD performed collection and processing of samples. AMH performed mass spectrometry-based procedures and analyses. MRB, AMH and JS performed all data analysis, statistical analysis and representation.

*Conceptualization: JS, HN, YC, SB*

*Data curation: MRB, AMH, JS, CD, FC, YC, SB*

*Formal analysis: MRB, AMH, JS, CD, FC, YC, SB*

*Funding acquisition HN, YC, SB*

*Investigation: MRB, AMH, JS, CD, HN, YC, SB*

*Methodology: AMH, HN, YC, SB*

*Project administration HN, YC, SB*

*Resources HN, YC, SB*

*Software*

*Supervision HN, YC, SB*

*Validation: AMH, JS, YC, SB*

*Visualization*

*Writing – original draft: MRB, AMH, JS, CD, HN, YC, SB*

*Writing – review & editing*

## SUPPORTING INFORMATION CAPTIONS

**Supplementary Figure 1:**
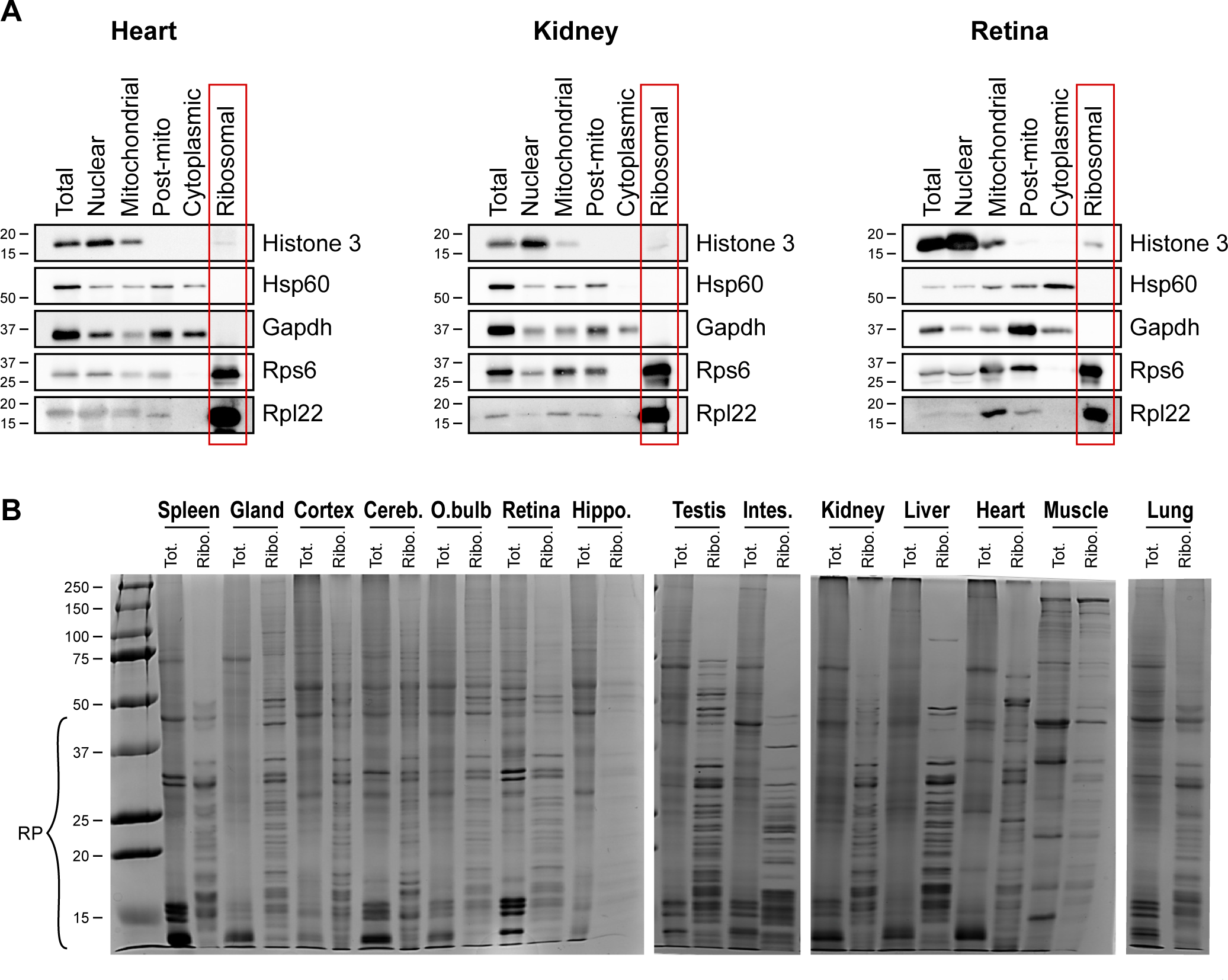
Validation of ribosomal fraction purification from adult mouse tissues. (A) Western blot analysis of markers of the different fractions obtained in the heart, the kidney and the retina: Histone 3 (nuclear), Hsp60 (mitochondrial and cytoplasmic), Gapdh (cytoplasmic), Rps6 and Rpl22 (ribosomal). (B) Protein profiles of the total and ribosomal fractions of each organ/tissue observed by Coomassie blue staining of proteins after SDS-PAGE. Molecular weights (kDa) are indicated on the left based on protein ladder.

**Supplementary Figure 2:**
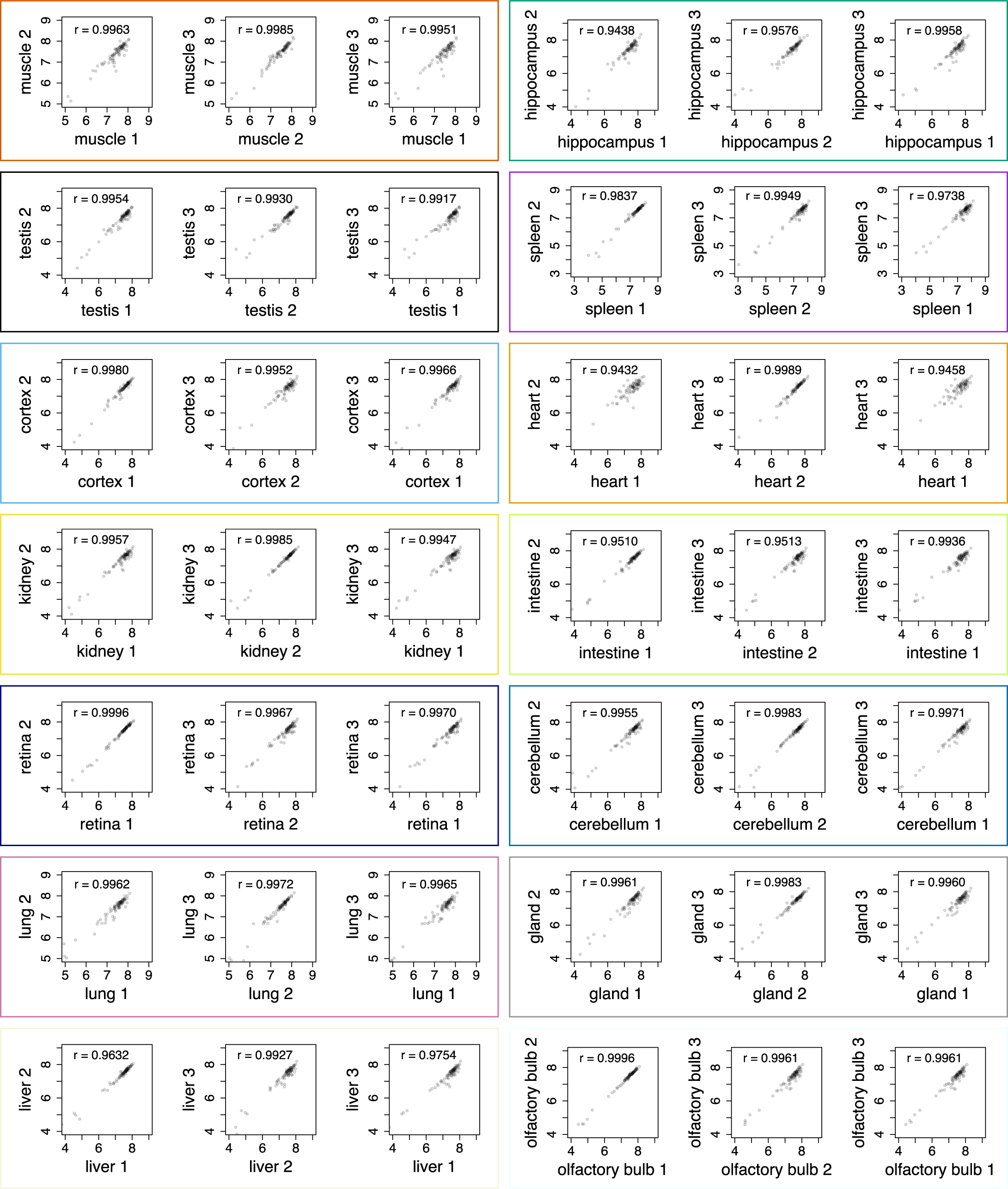
High consistency between biological replicates of the ribosomal fractions prepared from each tissue. Scatterplots of log-transformed protein abundance of the 85 RPs detected across replicates. The Pearson correlation coefficient is indicated on each plot.

**Supplementary Figure 3:**
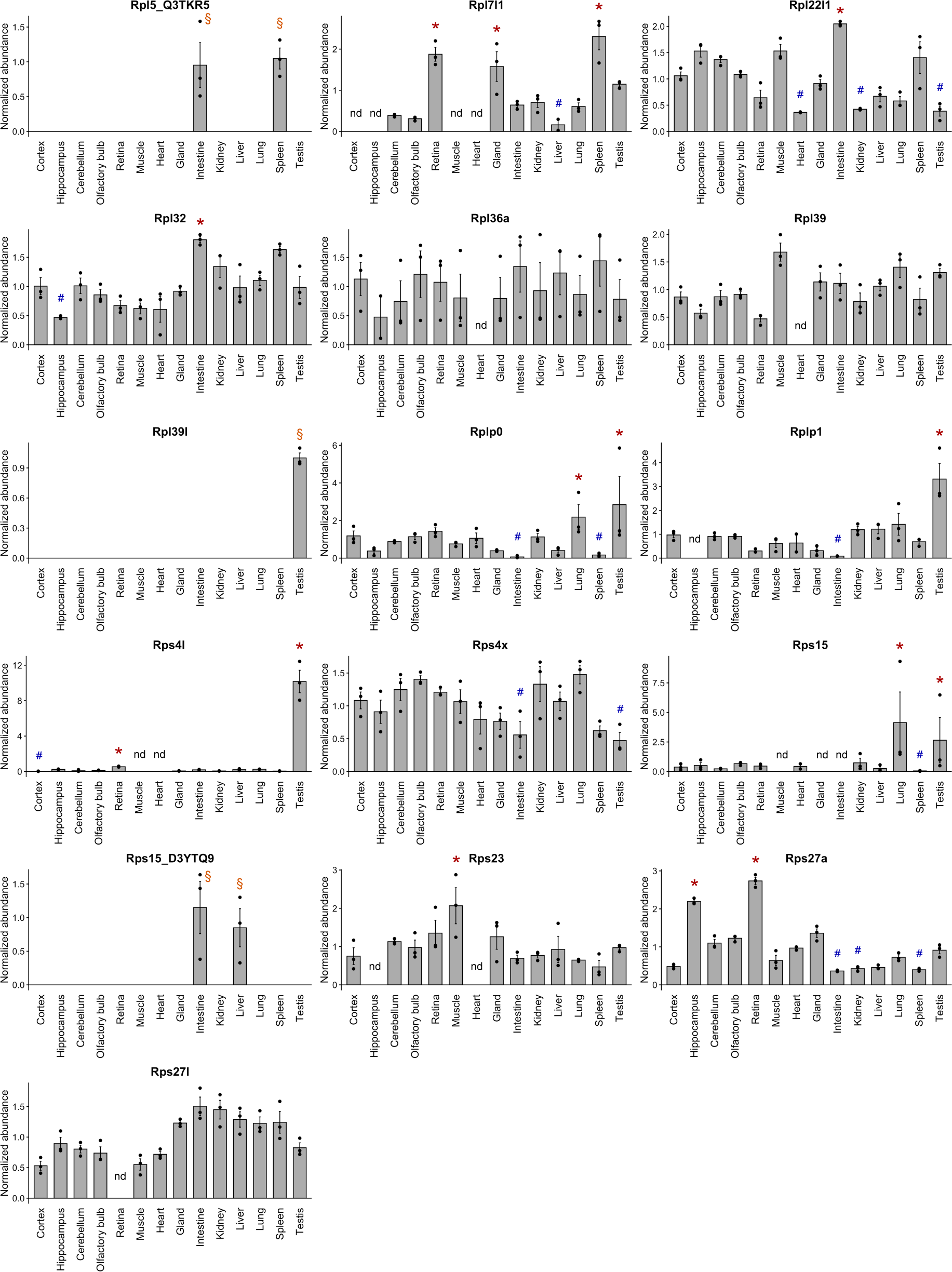
Variations in RP composition among adult mouse organs and tissues. Barplot representation of the relative abundance of variable RPs normalized to the sum of all RPs. The mean +/- s.e.m. is plotted for each organ, as well as values of individual replicates. * q-value < 0.01 (LIMMA test) and log_2_FC > 1 for One versus All comparisons. ^#^ q-value < 0.01 (LIMMA test) and log_2_FC < -1 for One versus All comparisons. ^§^ specific detection, q-value < 0.01 (ANOVA test). nd: not detected, q-value < 0.01 (LIMMA test) and no log_2_FC calculable for One versus All comparisons.

**Supplementary Figure 4:**
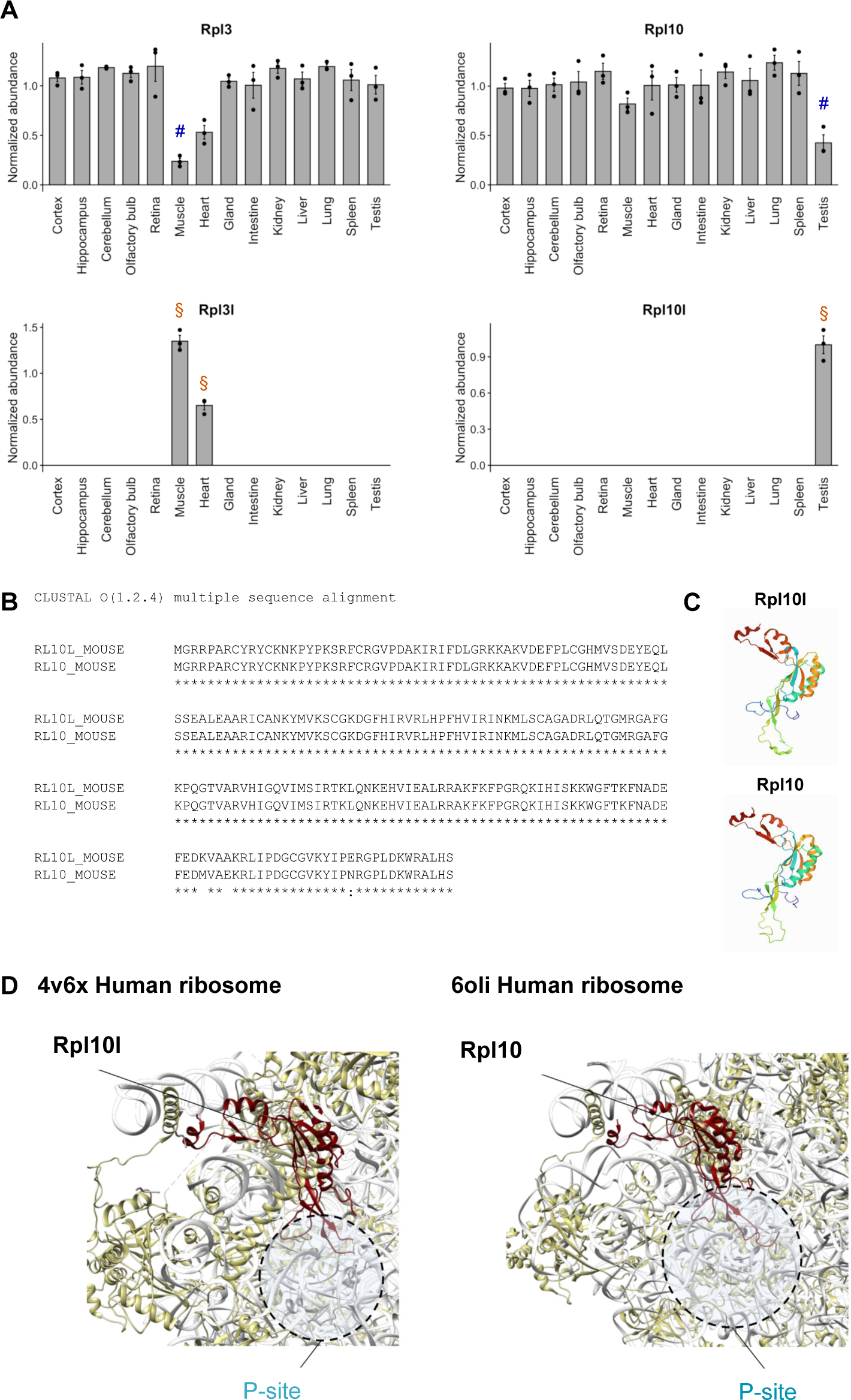
Paralogous RPs and corresponding canonical RPs show balanced enrichment in specific organs. (A) Barplot representation of the relative abundance of Rpl3l, Rpl3, Rpl10l and Rpl10 normalized to the sum of all RPs. The mean +/- s.e.m. is plotted for each organ, as well as values of individual replicates. ^#^ q-value < 0.01 (LIMMA test) and log_2_FC < -1 for One versus All comparisons. ^§^ specific detection, q-value < 0.01 (ANOVA test). (B) Alignment of amino acid sequences of mouse Rpl10l and mouse Rpl10. (C) 3D schematic representation of the molecular structure of Rpl10l and Rpl10. (D) Visualization of the localization of Rpl10l and Rpl10 in the molecular structure of the ribosome.

**Supplementary Figure 5:**
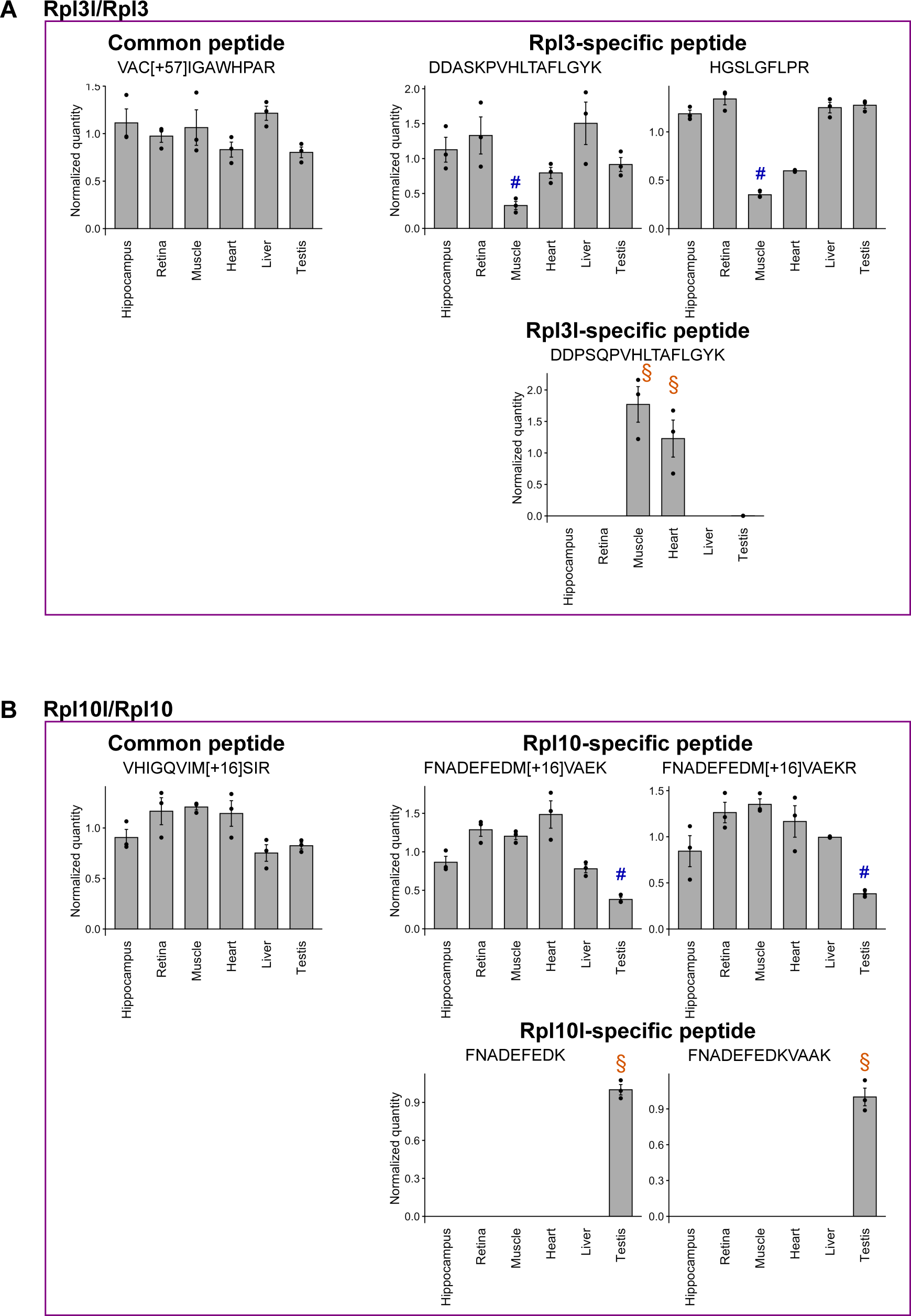
Targeted proteomics validates differential abundance of corresponding paralogous and canonical RPs in ribosomal fractions of adult mouse tissues. (A) Barplot representation of the relative abundance in selected tissues of peptides shared by specific to paralogous Rpl3l and to its corresponding canonical form Rpl3. (B) Barplot representation of the relative abundance in selected tissues of peptides shared by or specific to paralogous Rpl10l and to its corresponding canonical form Rpl10. The mean +/- s.e.m. is plotted for each organ, as well as values of individual replicates. ^#^q-value < 0.01 (LIMMA test) and log_2_FC < -1 for One versus All comparisons. ^§^ specific detection.

**Supplementary Figure 6:**
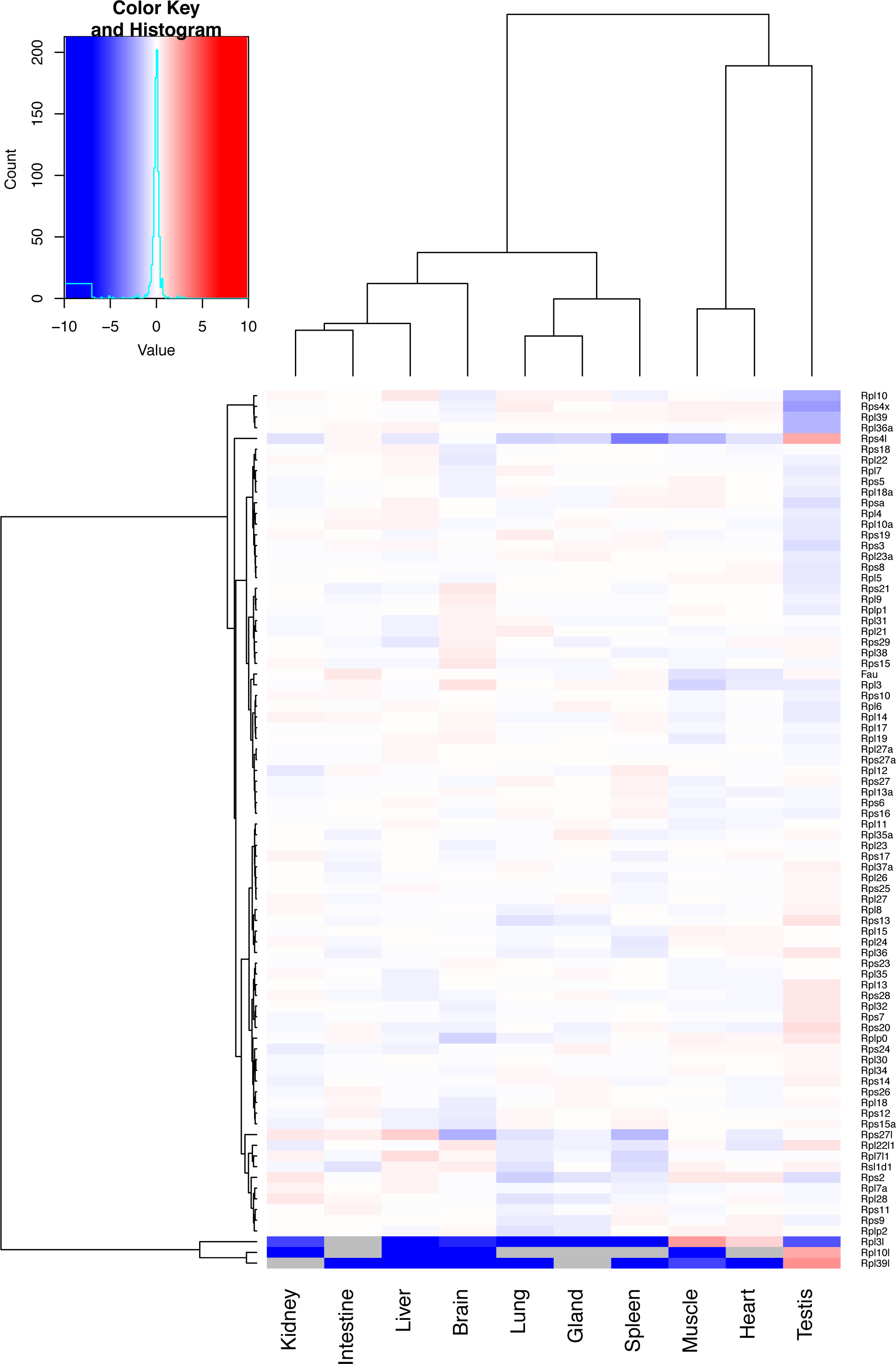
Relative expression of RP transcripts across adult mouse tissues. Heatmap of the log-transformed relative expression of RPs detected by RNA-sequencing, from depleted in blue to enriched in red. Grey boxes represent no detection of the RP transcript in the corresponding organ. Expression values from the Mouse Transcriptomic Body Map dataset (57).

**Supplementary Table 1: List of proteins detected in the ribosomal fraction of each adult mouse tissue and corresponding iBAQ values.**

**Supplementary Table 2: Results from MS-based label-free quantitative proteomic analyses. Details of ANOVA results.**

**Supplementary Table 3: Results from MS-based label-free quantitative proteomic analyses. Details of One versus All LIMMA test.**

**Supplementary Table 4: Quantification of selected peptides using targeted proteomics. Measured amounts are indicated in amol.**

**Supplementary Table 5: Results of ANOVA test and One versus All LIMMA test for targeted proteomics.**

## REFERENCES

1. Lee TI, Young RA. Transcriptional Regulation and its Misregulation in Disease. Cell. 2013 Mar 14;152(6):1237–51.

2. Casamassimi A, Ciccodicola A. Transcriptional Regulation: Molecules, Involved Mechanisms, and Misregulation. Int J Mol Sci. 2019 Mar 14;20(6):1281.

3. Jaenisch R, Bird A. Epigenetic regulation of gene expression: how the genome integrates intrinsic and environmental signals. Nat Genet. 2003 Mar;33(3):245–54.

4. Tran NM, Shekhar K, Whitney IE, Jacobi A, Benhar I, Hong G, et al. Single-Cell Profiles of Retinal Ganglion Cells Differing in Resilience to Injury Reveal Neuroprotective Genes. Neuron. 2019 Dec 18;104(6):1039–1055.e12.

5. Winter CC, Jacobi A, Su J, Chung L, van Velthoven CTJ, Yao Z, et al. A transcriptomic taxonomy of mouse brain-wide spinal projecting neurons. Nature. 2023 Dec;624(7991):403–14.

6. Casamassimi A, Federico A, Rienzo M, Esposito S, Ciccodicola A. Transcriptome Profiling in Human Diseases: New Advances and Perspectives. Int J Mol Sci. 2017 Jul 29;18(8):1652.

7. Mathys H, Davila-Velderrain J, Peng Z, Gao F, Mohammadi S, Young JZ, et al. Single-cell transcriptomic analysis of Alzheimer’s disease. Nature. 2019 Jun;570(7761):332–7.

8. Liu Y, Beyer A, Aebersold R. On the Dependency of Cellular Protein Levels on mRNA Abundance. Cell. 2016 Apr 21;165(3):535–50.

9. Fortelny N, Overall CM, Pavlidis P, Freue GVC. Can we predict protein from mRNA levels? Nature. 2017 Jul;547(7664):E19–20.

10. Schwanhäusser B, Busse D, Li N, Dittmar G, Schuchhardt J, Wolf J, et al. Global quantification of mammalian gene expression control. Nature. 2011 May;473(7347):337–42.

11. Vogel C, Marcotte EM. Insights into the regulation of protein abundance from proteomic and transcriptomic analyses. Nat Rev Genet. 2012 Mar 13;13(4):227–32.

12. Wilhelm M, Schlegl J, Hahne H, Gholami AM, Lieberenz M, Savitski MM, et al. Mass-spectrometry-based draft of the human proteome. Nature. 2014 May;509(7502):582–7.

13. Liu Y, Borel C, Li L, Müller T, Williams EG, Germain PL, et al. Systematic proteome and proteostasis profiling in human Trisomy 21 fibroblast cells. Nat Commun. 2017 Oct 31;8:1212.

14. Wang D, Eraslan B, Wieland T, Hallström B, Hopf T, Zolg DP, et al. A deep proteome and transcriptome abundance atlas of 29 healthy human tissues. Molecular Systems Biology. 2019 Feb;15(2):e8503.

15. Hamm DC, Paatela EM, Bennett SR, Wong CJ, Campbell AE, Wladyka CL, et al. The transcription factor DUX4 orchestrates translational reprogramming by broadly suppressing translation efficiency and promoting expression of DUX4-induced mRNAs. PLoS Biol. 2023 Sep;21(9):e3002317.

16. Li YF, Cheng T, Zhang YJ, Fu XX, Mo J, Zhao GQ, et al. Mycn regulates intestinal development through ribosomal biogenesis in a zebrafish model of Feingold syndrome 1. PLoS Biol. 2022 Nov;20(11):e3001856.

17. Brunet MA, Lucier JF, Levesque M, Leblanc S, Jacques JF, Al-Saedi HRH, et al. OpenProt 2021: deeper functional annotation of the coding potential of eukaryotic genomes. Nucleic Acids Res. 2020 Nov 12;49(D1):D380–8.

18. Cardon T, Fournier I, Salzet M. Shedding Light on the Ghost Proteome. Trends in Biochemical Sciences. 2021 Mar 1;46(3):239–50.

19. Petrov AS, Bernier CR, Hsiao C, Norris AM, Kovacs NA, Waterbury CC, et al. Evolution of the ribosome at atomic resolution. Proc Natl Acad Sci U S A. 2014 Jul 15;111(28):10251–6.

20. Roberts E, Sethi A, Montoya J, Woese CR, Luthey-Schulten Z. Molecular signatures of ribosomal evolution. Proc Natl Acad Sci U S A. 2008 Sep 16;105(37):13953–8.

21. Cech TR. The Ribosome Is a Ribozyme. Science. 2000 Aug 11;289(5481):878–9.

22. Genuth NR, Barna M. The discovery of ribosome heterogeneity and its implications for gene regulation and organismal life. Mol Cell. 2018 Aug 2;71(3):364–74.

23. Gay DM, Lund AH, Jansson MD. Translational control through ribosome heterogeneity and functional specialization. Trends in Biochemical Sciences [Internet]. 2021 Jul 23 [cited 2021 Jul 26];0(0). Available from: https://www.cell.com/trends/biochemical-sciences/abstract/S0968-0004(21)00144-4

24. Ulirsch JC, Verboon JM, Kazerounian S, Guo MH, Yuan D, Ludwig LS, et al. The Genetic Landscape of Diamond-Blackfan Anemia. The American Journal of Human Genetics. 2018 Dec 6;103(6):930–47.

25. Da Costa LM, Marie I, Leblanc TM. Diamond-Blackfan anemia. Hematology Am Soc Hematol Educ Program. 2021 Dec 10;2021(1):353–60.

26. Girardi T, Vereecke S, Sulima SO, Khan Y, Fancello L, Briggs JW, et al. The T-cell leukemia associated ribosomal RPL10 R98S mutation enhances JAK-STAT signaling. Leukemia. 2018 Mar;32(3):809–19.

27. Kampen KR, Sulima SO, Verbelen B, Girardi T, Vereecke S, Rinaldi G, et al. The ribosomal RPL10 R98S mutation drives IRES-dependent BCL-2 translation in T-ALL. Leukemia. 2019 Feb;33(2):319–32.

28. Ebert BL, Pretz J, Bosco J, Chang CY, Tamayo P, Galili N, et al. Identification of RPS14 as a 5q-syndrome gene by RNA interference screen. Nature. 2008 Jan 17;451(7176):335–9.

29. Sulima SO, Patchett S, Advani VM, De Keersmaecker K, Johnson AW, Dinman JD. Bypass of the pre-60S ribosomal quality control as a pathway to oncogenesis. Proc Natl Acad Sci U S A. 2014 Apr 15;111(15):5640–5.

30. Lezzerini M, Penzo M, O’Donohue MF, Marques dos Santos Vieira C, Saby M, Elfrink HL, et al. Ribosomal protein gene RPL9 variants can differentially impair ribosome function and cellular metabolism. Nucleic Acids Res. 2020 Jan 24;48(2):770–87.

31. Decourt C, Schaeffer J, Blot B, Paccard A, Excoffier B, Pende M, et al. The RSK2-RPS6 axis promotes axonal regeneration in the peripheral and central nervous systems. PLOS Biology. 2023 Apr 17;21(4):e3002044.

32. Chaillou T, Zhang X, McCarthy JJ. Expression of muscle-specific ribosomal protein L3-like impairs myotube growth. J Cell Physiol. 2016 Sep;231(9):1894–902.

33. Jiang L, Li T, Zhang X, Zhang B, Yu C, Li Y, et al. RPL10L Is Required for Male Meiotic Division by Compensating for RPL10 during Meiotic Sex Chromosome Inactivation in Mice. Curr Biol. 2017 May 22;27(10):1498–1505.e6.

34. Komili S, Farny NG, Roth FP, Silver PA. Functional specificity among ribosomal proteins regulates gene expression. Cell. 2007 Nov 2;131(3):557–71.

35. Shi Z, Fujii K, Kovary KM, Genuth NR, Röst HL, Teruel MN, et al. Heterogeneous Ribosomes Preferentially Translate Distinct Subpools of mRNAs Genome-wide. Molecular Cell. 2017 Jul 6;67(1):71–83.e7.

36. Lee ASY, Burdeinick-Kerr R, Whelan SPJ. A ribosome-specialized translation initiation pathway is required for cap-dependent translation of vesicular stomatitis virus mRNAs. Proc Natl Acad Sci U S A. 2013 Jan 2;110(1):324–9.

37. Kondrashov N, Pusic A, Stumpf CR, Shimizu K, Hsieh AC, Xue S, et al. Ribosome-Mediated Specificity in Hox mRNA Translation and Vertebrate Tissue Patterning. Cell. 2011 Apr 29;145(3):383–97.

38. Xue S, Tian S, Fujii K, Kladwang W, Das R, Barna M. RNA regulons in Hox 5′UTRs confer ribosome specificity to gene regulation. Nature. 2015 Jan 1;517(7532):33–8.

39. Ivanov IP, Saba JA, Fan CM, Wang J, Firth AE, Cao C, et al. Evolutionarily conserved inhibitory uORFs sensitize Hox mRNA translation to start codon selection stringency. Proc Natl Acad Sci U S A. 2022 Mar 1;119(9):e2117226119.

40. Slavov N, Semrau S, Airoldi E, Budnik B, van Oudenaarden A. Differential Stoichiometry among Core Ribosomal Proteins. Cell Reports. 2015 Nov 3;13(5):865–73.

41. Petelski AA, Slavov N. Analyzing Ribosome Remodeling in Health and Disease. Proteomics. 2020 Sep;20(17–18):e2000039.

42. Yoon A, Peng G, Brandenburg Y, Zollo O, Xu W, Rego E, et al. Impaired Control of IRES-Mediated Translation in X-Linked Dyskeratosis Congenita. Science. 2006 May 12;312(5775):902–6.

43. Belin S, Beghin A, Solano-Gonzàlez E, Bezin L, Brunet-Manquat S, Textoris J, et al. Dysregulation of Ribosome Biogenesis and Translational Capacity Is Associated with Tumor Progression of Human Breast Cancer Cells. PLoS One. 2009 Sep 25;4(9):e7147.

44. Marcel V, Ghayad SE, Belin S, Therizols G, Morel AP, Solano-Gonzàlez E, et al. p53 Acts as a Safeguard of Translational Control by Regulating Fibrillarin and rRNA Methylation in Cancer. Cancer Cell. 2013 Sep 9;24(3):318–30.

45. Erales J, Marchand V, Panthu B, Gillot S, Belin S, Ghayad SE, et al. Evidence for rRNA 2′-O-methylation plasticity: Control of intrinsic translational capabilities of human ribosomes. Proc Natl Acad Sci U S A. 2017 Dec 5;114(49):12934–9.

46. Leppek K, Fujii K, Quade N, Susanto TT, Boehringer D, Lenarčič T, et al. Gene- and Species-Specific Hox mRNA Translation by Ribosome Expansion Segments. Molecular Cell [Internet]. 2020 Nov 16 [cited 2020 Nov 23]; Available from: http://www.sciencedirect.com/science/article/pii/S1097276520307309

47. Akirtava C, May GE, McManus CJ. False-positive IRESes from Hoxa9 and other genes resulting from errors in mammalian 5′ UTR annotations. Proc Natl Acad Sci U S A. 2022 Sep 6;119(36):e2122170119.

48. Amirbeigiarab S, Kiani P, Sanchez AV, Krisp C, Kazantsev A, Fester L, et al. Invariable stoichiometry of ribosomal proteins in mouse brain tissues with aging. PNAS. 2019 Nov 5;116(45):22567–72.

49. Gupta V, Warner JR. Ribosome-omics of the human ribosome. RNA. 2014 Jul;20(7):1004–13.

50. Guimaraes JC, Zavolan M. Patterns of ribosomal protein expression specify normal and malignant human cells. Genome Biology. 2016 Nov 24;17(1):236.

51. Li H, Huo Y, He X, Yao L, Zhang H, Cui Y, et al. A male germ-cell-specific ribosome controls male fertility. Nature. 2022 Dec;612(7941):725–31.

52. Alkan F, Wilkins OG, Hernández-Pérez S, Ramalho S, Silva J, Ule J, et al. Identifying ribosome heterogeneity using ribosome profiling. Nucleic Acids Res. 2022 Jun 10;50(16):e95.

53. Belin S, Hacot S, Daudignon L, Therizols G, Pourpe S, Mertani HC, et al. Purification of Ribosomes from Human Cell Lines. Current Protocols in Cell Biology. 2010;49(1):3.40.1-3.40.11.

54. Wool IG, Chan YL, Glück A. Structure and evolution of mammalian ribosomal proteins. Biochem Cell Biol. 1995 Dec;73(11–12):933–47.

55. Hopes T, Norris K, Agapiou M, McCarthy CGP, Lewis PA, O’Connell MJ, et al. Ribosome heterogeneity in Drosophila melanogaster gonads through paralog-switching. Nucleic Acids Res. 2021 Jul 20;50(4):2240–57.

56. Mageeney CM, Ware VC. Specialized eRpL22 paralogue-specific ribosomes regulate specific mRNA translation in spermatogenesis in Drosophila melanogaster. Mol Biol Cell. 2019 Aug 1;30(17):2240–53.

57. Li B, Qing T, Zhu J, Wen Z, Yu Y, Fukumura R, et al. A Comprehensive Mouse Transcriptomic BodyMap across 17 Tissues by RNA-seq. Sci Rep. 2017 Jun 23;7(1):4200.

58. Yue F, Cheng Y, Breschi A, Vierstra J, Wu W, Ryba T, et al. A comparative encyclopedia of DNA elements in the mouse genome. Nature. 2014 Nov 20;515(7527):355–64.

59. Kampen KR, Sulima SO, De Keersmaecker K. Rise of the specialized onco-ribosomes. Oncotarget. 2018 Oct 16;9(81):35205–6.

60. Simsek D, Tiu GC, Flynn RA, Byeon GW, Leppek K, Xu AF, et al. The Mammalian Ribo-interactome Reveals Ribosome Functional Diversity and Heterogeneity. Cell. 2017 Jun;169(6):1051–1065.e18.

61. Xue S, Barna M. Specialized ribosomes: a new frontier in gene regulation and organismal biology. Nat Rev Mol Cell Biol. 2012 May 23;13(6):355–69.

62. Gemmer M, Chaillet ML, van Loenhout J, Cuevas Arenas R, Vismpas D, Gröllers-Mulderij M, et al. Visualization of translation and protein biogenesis at the ER membrane. Nature. 2023 Feb;614(7946):160–7.

63. O’Leary MN, Schreiber KH, Zhang Y, Duc ACE, Rao S, Hale JS, et al. The Ribosomal Protein Rpl22 Controls Ribosome Composition by Directly Repressing Expression of Its Own Paralog, Rpl22l1. PLOS Genetics. 2013 Aug 22;9(8):e1003708.

64. Shiraishi C, Matsumoto A, Ichihara K, Yamamoto T, Yokoyama T, Mizoo T, et al. RPL3L-containing ribosomes determine translation elongation dynamics required for cardiac function. Nat Commun. 2023 Apr 20;14(1):2131.

65. Mageeney CM, Kearse MG, Gershman BW, Pritchard CE, Colquhoun JM, Ware VC. Functional interplay between ribosomal protein paralogues in the eRpL22 family in Drosophila melanogaster. Fly (Austin). 2018 Nov 29;12(3–4):143–63.

66. Segev N, Gerst JE. Specialized ribosomes and specific ribosomal protein paralogs control translation of mitochondrial proteins. J Cell Biol. 2018 Jan 2;217(1):117–26.

67. Ghulam MM, Catala M, Abou Elela S. Differential expression of duplicated ribosomal protein genes modifies ribosome composition in response to stress. Nucleic Acids Res. 2020 Feb 28;48(4):1954–68.

68. Petibon C, Ghulam MM, Catala M, Elela SA. Regulation of ribosomal protein genes: An ordered anarchy. WIREs RNA. 2021;12(3):e1632.

69. Panda A, Yadav A, Yeerna H, Singh A, Biehl M, Lux M, et al. Tissue- and development-stage–specific mRNA and heterogeneous CNV signatures of human ribosomal proteins in normal and cancer samples. Nucleic Acids Res. 2020 Jul 27;48(13):7079–98.

70. Warner JR, McIntosh KB. How Common are Extra-ribosomal Functions of Ribosomal Proteins? Mol Cell. 2009 Apr 10;34(1):3–11.

71. Ishii K, Washio T, Uechi T, Yoshihama M, Kenmochi N, Tomita M. Characteristics and clustering of human ribosomal protein genes. BMC Genomics. 2006 Feb 28;7(1):37.

72. Ballesta JP, Remacha M. The large ribosomal subunit stalk as a regulatory element of the eukaryotic translational machinery. Prog Nucleic Acid Res Mol Biol. 1996;55:157–93.

73. Shigeoka T, Koppers M, Wong HHW, Lin JQ, Cagnetta R, Dwivedy A, et al. On-Site Ribosome Remodeling by Locally Synthesized Ribosomal Proteins in Axons. Cell Reports. 2019 Dec 10;29(11):3605–3619.e10.

74. Fusco CM, Desch K, Dörrbaum AR, Wang M, Staab A, Chan ICW, et al. Neuronal ribosomes exhibit dynamic and context-dependent exchange of ribosomal proteins. Nat Commun. 2021 Oct 21;12(1):6127.

75. Salvetti A, Couté Y, Epstein A, Arata L, Kraut A, Navratil V, et al. Nuclear Functions of Nucleolin through Global Proteomics and Interactomic Approaches. J Proteome Res. 2016 May 6;15(5):1659–69.

76. Bouyssié D, Hesse AM, Mouton-Barbosa E, Rompais M, Macron C, Carapito C, et al. Proline: an efficient and user-friendly software suite for large-scale proteomics. Bioinformatics. 2020 May 15;36(10):3148–55.

77. Wieczorek S, Combes F, Lazar C, Giai Gianetto Q, Gatto L, Dorffer A, et al. DAPAR & ProStaR: software to perform statistical analyses in quantitative discovery proteomics. Bioinformatics. 2017 Jan 1;33(1):135–6.

78. R Core Team. R: A language and environment for statistical computing. Vienna, Austria: R Foundation for Statistical Computing; 2014. 2014.

79. Pettersen EF, Goddard TD, Huang CC, Couch GS, Greenblatt DM, Meng EC, et al. UCSF Chimera--a visualization system for exploratory research and analysis. J Comput Chem. 2004 Oct;25(13):1605–12.

80. Perez-Riverol Y, Csordas A, Bai J, Bernal-Llinares M, Hewapathirana S, Kundu DJ, et al. The PRIDE database and related tools and resources in 2019: improving support for quantification data. Nucleic Acids Res. 2019 Jan 8;47(Database issue):D442–50.

